# Plasmid fitness costs are caused by specific genetic conflicts

**DOI:** 10.1101/2021.04.10.439128

**Authors:** James P. J. Hall, Rosanna C. T. Wright, Ellie Harrison, Katie J. Muddiman, A. Jamie Wood, Steve Paterson, Michael A. Brockhurst

**Affiliations:** Department of Evolution, Ecology and Behaviour, Institute of Infection, Veterinary and Ecological Sciences, University of Liverpool, Crown Street, Liverpool, L69 7ZB; Department of Animal and Plant Sciences, University of Sheffield, Western Bank, Sheffield, S10 2TN; Division of Evolution and Genomic Sciences, University of Manchester, Oxford Road, Manchester, M13 9PL; Department of Biology, University of York, Wentworth Way, York, YO10 5DD; Department of Mathematics, University of York, Wentworth Way, York, YO10 5DD

## Abstract

Plasmids play an important role in bacterial genome evolution by transferring genes between lineages. Fitness costs associated with plasmid acquisition are expected to be a barrier to gene exchange, but the causes of plasmid fitness costs are poorly understood. Single compensatory mutations are often sufficient to completely ameliorate plasmid fitness costs, suggesting that such costs are caused by specific genetic conflicts rather than generic properties of plasmids, such as their size, metabolic burden, or expression level. Here we show — using a combination of experimental evolution, reverse genetics, and transcriptomics — that fitness costs of two divergent large plasmids in *Pseudomonas fluorescens* are caused by inducing maladaptive expression of a chromosomal tailocin toxin operon. Mutations in single genes unrelated to the toxin operon, and located on either the chromosome or the plasmid, ameliorated the disruption associated with plasmid acquisition. We identify one of these compensatory loci, the chromosomal gene *PFLU4242*, as the key mediator of the fitness costs of both plasmids, with the other compensatory loci either reducing expression of this gene or mitigating its deleterious effects by upregulating a putative plasmid-borne ParAB operon. The chromosomal mobile genetic element Tn6291, which uses plasmids for transmission, remained upregulated even in compensated strains, suggesting that mobile genetic elements communicate through pathways independent of general physiological disruption. Plasmid fitness costs caused by specific genetic conflicts are unlikely to act as a long-term barrier to horizontal gene transfer due to their propensity for amelioration by single compensatory mutations, explaining why plasmids are so common in bacterial genomes.

## Introduction

Plasmid-mediated horizontal gene transfer (HGT) is a powerful force in bacterial evolution. Many of the traits necessary for the ecological success of bacteria in new environments are carried between bacterial strains and species by plasmids. In hospitals, plasmids mobilise antimicrobial resistance genes between pathogens, commensals, and environmental bacteria of diverse species [1–3]. In soil, plasmids are responsible for bioremediation of contaminated sites, and provide new ecological opportunities both to free-living and to plant-associated bacteria [4, 5]. In infections, plasmids provide key virulence determinants necessary for colonisation and establishment, and factors that assist in immune evasion and pathogen transmission [6, 7]. Thus in diverse habitats, plasmids are integral constituents of microbiomes, acting as potent drivers of bacterial evolutionary ecology, in large part through their role as vehicles of HGT [8–11].

Nevertheless, there exist various features of their bacterial hosts that restrict plasmid transmission, and thus the flow of genetic information [12]. Some of these features impede acquisition of plasmids, such as entry exclusion systems, CRISPR, and restriction systems [13, 14]. Other barriers emerge from the inability of plasmids to replicate or to segregate effectively to daughter cells. However, even if a plasmid can enter, replicate, and be passed to daughter cells, its longer term persistence is contingent upon the fitness costs that emerge from plasmid carriage [15, 16].

Plasmid-imposed fitness costs are the negative impacts on growth, replication, and/or survival that emerge as a consequence of plasmid carriage [15]. These costs are thought to act as major barriers to plasmid maintenance — and plasmid-mediated HGT — because cells burdened by the costs of plasmid carriage are outcompeted by others that lack the plasmid and its associated costs [17]. Importantly, selection for plasmid-borne genes (for example, by antibiotics) is insufficient to explain plasmid persistence in the long term, because those genes can become relocated to the chromosome through recombination, resulting in net fitness costs of plasmid carriage, favouring plasmid loss [17–19]. Therefore, to understand plasmid maintenance — and by extension the ability of plasmids to act as vehicles of gene exchange in bacterial communities — it is crucial to understand the mechanisms that underlie plasmid fitness costs, and how these costs may be resolved.

A variety of causes of plasmid fitness costs have been proposed [15], but in most cases the underlying molecular mechanisms remain unknown. General properties of plasmids may explain costs because cellular resources — not least nucleotides, amino acids, and the machineries of replication, transcription, and translation — must be redirected to plasmid activity, imposing a metabolic burden [16, 20]. These costs would be expected to scale with plasmid size, copy number, and gene expression, being higher for plasmids that are large, multi-copy, and/or have highly active promoters. Plasmid DNA composition, particularly GC content and bias in codon use, is another general property that can be a potential source of plasmid costs, exacerbating the inefficiencies in bacterial physiology caused by plasmid maintenance [21, 22]. Alternatively, plasmid fitness costs can derive from the effects of specific plasmid genes. Some plasmid-borne genes are costly in and of themselves, for example some efflux pumps [23] or the conjugative pilus which can disrupt the cell membrane and exposes bacteria to phage predation [24], whereas other plasmid genes elicit deleterious interactions with chromosomal genes, for example by triggering stress responses [25] or interfering with metabolism [26, 27].

Commonly, plasmids or the host bacterium undergo compensatory evolution, reducing the costs of plasmid carriage and hence the strength of purifying selection against plasmids, enabling maintenance [15,23,28–33]. Intuitively, the nature of compensatory mutations can provide an insight into the causes of plasmid fitness costs. If costs emerge from general properties of plasmids, we might expect large scale changes, such as deletions that reduce plasmid size, or many individual changes to plasmid codon usage. Whereas, if costs derive from specific plasmid genes or specific plasmid-chromosomal gene interactions, we might instead expect targeted deletions, mutations, or regulatory changes affecting only those genes. Whilst some studies of compensatory evolution do report large-scale plasmid deletions [28, 34], the majority implicate one or few mutations to specific genes as responsible for ameliorating the cost of plasmid carriage [23,25,29–33], suggesting that specific genetic interactions may commonly underlie plasmid fitness costs. Identifying these interactions, and how they are resolved by compensatory evolution, is crucial for understanding the evolution and ecology of horizontal gene transfer. In particular, the location of compensatory evolution is predicted to have important consequences: if the compensatory mutation occurs to the plasmid then it is propagated with onward horizontal transmission [30, 35], whereas mutations occurring on the chromosome can potentially facilitate accumulation of many different plasmids [32, 36]. However, chromosomal and plasmid compensations may not both be available for some plasmid-host combinations owing to the processes by which plasmid costs emerge and are resolved. Although many compensatory mutations have been identified for different plasmid-bacterial pairings, the mechanistic bases of compensation, and thus the nature of the resolved fitness costs, often remain unclear.

Here, we take advantage of a plasmid-host system where multiple genetic pathways of compensatory evolution exist, to investigate the mechanistic bases of plasmid fitness costs and their amelioration. We have previously described two pathways of chromosomal compensatory mutation in *Pseudomonas fluorescens* SBW25, targeting the *gacA/S* global regulatory system or the hypothetical protein *PFLU4242*, which individually ameliorate the fitness costs of megaplasmids pQBR103 (425 kb), pQBR57 (307 kb), or pQBR55 (157 kb) [29, 37]. In this study, we describe an additional plasmid-borne compensatory locus, the lambda-repressor-like protein PQBR57_0059, that ameliorates the cost of pQBR57. Using a combination of experimental evolution, transcriptomics and reverse genetics we then show that specific genetic conflicts between chromosomal and plasmid genes, not plasmid size or gene expression *per se*, are the principal cause of plasmid fitness costs in this system, and identify a single chromosomal locus that acts as the key mediator of plasmid fitness costs.

## Materials and methods

### Bacterial strains

*Pseudomonas fluorescens* SBW25 was cultured in 6 ml King’s B (KB) media in 30 ml glass universals (‘microcosms’) at 28°C shaking at 180 rpm. Strains were labelled with either gentamicin resistance (GmR) or streptomycin resistance (SmR) by insertion of mini-Tn7 resistance cassettes into *attTn7* sites according to the protocol of Choi & Schweizer [38]. Allelic replacement of *gacA*, *gacS* and *PFLU4242* genes was achieved using the pUIC3 suicide vector with a two-step protocol as described previously [33]. The ‘evolved’ pQBR57 plasmids came from strains A.01.65.G.002 (ancestral type) and B.09.65.G.029 (V100A variant; Samples 1 and 49 respectively from [39]). These plasmids were allowed to transfer into an ancestral SmR background which were used as donor strains for conjugation into the experimental lines. Plasmid pQBR103 was from [40].

Transconjugants for use in experiments were generated by mixing equal volumes of overnight cultures of donors and recipients and co-culturing for 24 hours before spreading on KB plates containing mercuric chloride (20 µM) and either gentamicin (30 µg/ml) or streptomycin (250 µg/ml). Candidate transconjugant colonies were re-streaked on selective media and tested by PCR for the presence of the plasmid. Independent transconjugants were used for each replicate.

To express PQBR57_0059 independently of pQBR57, amplicons from the ‘anc’ and ‘V100A’ strains were cloned EcoRI/KpnI into pUCP18 [41]. Inserts were sequenced, and 500 ng plasmid was transformed into *P. fluorescens* SBW25 GmR using the sucrose method [38]. Three independent transformants were generated (one for each replicate). Putative pQBR57 *par* genes were cloned into pUCP18 and transformed into *P. fluorescens* SBW25 in a similar manner. pUCP18-containing lines were selected and maintained on kanamycin 50 µg/ml, and pQBR57 and pQBR103 variants were transferred to these lineages as described above. To test effects of expressing plasmid-regulated genes, candidates were cloned EcoRI/KpnI into pME6032 [42], sequenced, and transformed into *P. fluorescens* as above. pME6032-containing lines were selected and maintained on 10 µg/ml (*E. coli*) or 100 µg/ml (*P. fluorescens*) tetracycline, and expression was induced with 100 µM IPTG.

### Competitions

For all competitions, GmR strains were competed against SmR-*lacZ* [43] strains. Overnight cultures of competing strains were washed and resuspended in KB, mixed in approximately equal ratios, and used to inoculate a KB microcosm at a 1:100 dilution. Samples of the competitors were spread on KB agar to estimate colony forming units at the beginning of the experiment and after 48 hours, and relative fitness was calculated as the ratio of Malthusian parameters W = ln(test_end_/test_start_)/ln(reference_end_/reference_start_) [44]. These assays are designed to assess the fate of lineages that initially start with the plasmid, rather than plasmids *per se*, since competitors may gain or lose the plasmid during the assay. Nevertheless, replica plating showed that segregation had a negligible effect on the overall proportions of plasmid-bearers over the 48 hours of the competition (<0.01% of initially resistant colonies became susceptible, see also Hall et al. [45]). For the highest conjugation rate conditions (into *P. fluorescens*, where the donor has both compensatory mutations), transconjugants comprised ∼10% of the final plasmid-bearing population. Measuring the fitness of the plasmid (rather than fitness of plasmid-bearers) increased calculated fitness measurements by <7% on average.

‘Medium-term’ competitions in soil were performed as described in [39]. Briefly, equal ratios of plasmid bearers and plasmid-free strains were mixed and added to soil microcosms, and transferred into fresh media every four days for 5 transfers in total. Populations were enumerated by plating or replica-plating onto selective media.

### Growth curves

Growth curves of pME6032-carrying lines were established from a 1:1000 dilution of overnight culture into 150 µl KB + tetracycline ± 100 µM IPTG. Plates with growth curve cultures were incubated under humid conditions at 28°C shaking at 180 rpm (3 mm oribital radius), and OD600 was measured every 15 minutes on a Tecan Spark 10M plate reader. Four independent replicate cultures were established for each of three independent transformants.

### RNA extraction and sequencing

RNA was extracted using TRI Reagent (T9424, Sigma Aldrich). Overnight cultures of bacteria were subcultured 1:50 into 6 ml prewarmed KB microcosms and grown at 28°C shaking at 180 rpm. When cultures reached intermediate exponential phase (OD600 ∼ 0.6, as assessed on a Tecan Spark plate reader) 1 ml samples were added to 0.4 volumes of ice-cold 95% ethanol 5% phenol ‘stop solution’ [46], mixed gently, and incubated on ice for at least 30 minutes. Each of the three replicates was processed separately as a block. Samples were harvested at 4.5 G for 10 minutes, resuspended in 1 ml TRI reagent, and incubated at room temperate for 5 minutes. Amylene-stabilised chloroform (400 µl, Sigma-Aldrich 34854-M) was added to each tube, and incubated for 2-15 minutes at room temperature. Samples were then transferred to 5Prime Phase Lock Gel ‘Heavy’ tubes (VWR 733-2478) and centrifuged at 17 G for 15 minutes at room temperature. The aqueous phase was collected into a fresh microfuge tube and 450 µl isopropanol was added to precipitate RNA. Samples were incubated at room temperature for 30 minutes before pelleting at 17 G for 30 minutes. The pellet was rinsed by gently adding 1 ml of 70% ethanol and pelleting at 7.5 G for 5 minutes. The supernatant was carefully removed and the pellet allowed to air dry before resuspension in RNase-free water at 65°C. To remove any residual contaminating DNA, RNA was diluted to ≤200 ng/µl and treated with TURBO DNA-free RNase-free DNase (Invitrogen AM1907) according to the manufacturer’s instructions. Reaction products were purified with Agencourt RNAClean XP beads. Ribosomal RNA was removed using the Bacterial RiboZero rRNA depletion kit (Illumina), and stranded libraries generated using NEBNext Ultra-Directional RNA library preparation kit (Illumina). Libraries were sequenced using paired-end 2×150 bp sequencing on two lanes of a HiSeq 4000. Short read sequences are available on NCBI-GEO short-read archive, project accession GSE151570.

### RNAseq analysis

Illumina adapter sequences were trimmed from the raw FASTQ files using Cutadapt version 1.2.1, using the option −O 3. The reads were further trimmed using Sickle version 1.200 (https://github.com/najoshi/sickle), with a minimum window quality score of 20, and reads shorter than 20 bp after trimming were removed. FastQC (http://www.bioinformatics.babraham.ac.uk/projects/fastqc/) and MultiQC [47] were used to analyse the reads. Warnings arising from “Sequence content,” “Sequence length” and “Sequence duplication” modules were followed up and found to be largely due to an overrepresentation of functional RNAs, namely tmRNA and RnaseP. Reads from each sample were aligned strand-specifically using HISAT2 version 2.1.0 [48] to the corresponding reference sequence(s): either the *P. fluorescens* SBW25 chromosome (AM181176) alone, the chromosome with pQBR57 (LN713926), or the chromosome with pQBR103 (AM235768). The options phred3, no-spliced-alignment, new-summary, and rna-strandness RF were used, and the output was filtered using samtools to remove mappings with PHRED-scaled quality < **A.** 10. The function featureCounts from the Rsubread package version 1.30.9 [49] was used to identify the reads mapping to each feature, and the table of counts was analysed in edgeR version 3.22.5 [50]. Here, we restricted differential expression analysis to putative protein-coding genes. Expression of chromosomal genes was analysed in a set of negative binomial GLMs and all treatments were compared with the plasmid-free ancestor with a series of quasi-likelihood F-tests. P-values were corrected using the Benjamini-Hochberg false-discovery rate of q = 0.05.

### Statistical analyses

Relative fitness of knockout strains was analysed in a linear mixed effects model using the R package nlme. The data were Box-Cox transformed to meet model assumptions. A model was fitted to all interactions of plasmid and host genotype. Transconjugant replicate was included as a random effect. Model reduction was performed using stepwise AIC comparisons and likelihood ratio tests. Effects of mutations and of plasmid carriage were estimated using post-hoc tests with p value adjustment using a multivariate *t* distribution using the lsmeans() function in the R package emmeans.

Relative fitness of plasmid mutants was analysed in a linear model with Box-Cox transformation. Though the effect of the *lacZ* marker was found to be non-significant (one-sample t-test, t_5_ = 0.32, p = 0.76), fitness values were initially corrected for marker effects by dividing each measurement by the mean fitness of plasmid-free strains (1.01). A linear model was performed with plasmid and marker and their interaction as fixed effects. Model reduction was performed using stepwise AIC comparisons and F tests. We detected no interaction effect of the marker (F_1,25_ = 0.03, p = 0.86). Post-hoc pairwise contrasts were performed as above.

Relative fitness of plasmid mutants was analysed in a linear model, with plasmid and knockout and their interactions as fixed effects, with post-hoc contrasts performed as above.

Relative fitness of strains expressing PQBR57_0059 *in trans*, or *par* in trans, were each analysed with linear mixed-effects models, with transformant as a random effect, and pUCP variant and pQBR variant and their interactions as fixed effects. Model reduction was performed using stepwise AIC comparisons and likelihood ratio tests, and post-hoc tests performed as above. The experiments with pQBR103 and *par* were analysed used linear models, as transconjugant replicates were not repeatedly-measured.

The dynamics of different PQBR57_0059 variants over time were analysed using a linear mixed-effects model. We used Box-Cox-transformed cumulative plasmid density as the response variable, and PQBR57_0059 status as a fixed effect. PQBR57_0059 variant was modelled using a random effect.

Growth curves were analysed using the ratios of cumulative densities of IPTG-induced and non-induced conditions as the response variable, and pME6032 insert as a fixed effect. Transformant replicate was modelled as a random effect. Post-hoc contrasts were performed for each strain against the no-insert control using the trt.vs.ctrl option in lsmeans(), with p value adjustment using the multivariate *t* distribution.

Data and example analysis scripts are available at https://github.com/jpjh/COMPMUT/ and at the University of Liverpool Datacat, doi: **10.17638/datacat.liverpool.ac.uk/1275**.

## Results

### Mutations to PFLU4242 and pQBR57_0059 emerged in parallel in evolution experiments

Previous evolution experiments with *P. fluorescens* SBW25 and megaplasmids pQBR103, pQBR57, or pQBR55, have identified four key genes implicated in plasmid compensatory evolution in this strain [29,33,37,39,51] (**Fig. 1**, full details and discussion in **Supplementary Text**): the *gacA/S* two-component signalling pathway, which has repeatedly been associated with pQBR103 and pQBR55 carriage; *PFLU4242*, a chromosomal gene that has been mutated in evolution experiments with all three pQBR plasmids tested; and *PQBR57_0059*, the only plasmid-borne gene that was a target of parallel evolution in pQBR57 [52]. Parallel mutations are a strong signal of selection [53, 54], and the appearance of similar mutations in experiments with different plasmids led us to hypothesise that mutations to *PFLU4242* and *gacS* were general mechanisms plasmid compensation in *P. fluorescens* SBW25. Given the difficulties of isolating wild-type pQBR55 transconjugants (i.e. without *de novo* compensation [29]), we focused on identifying the basis of fitness cost and amelioration in the the two larger megaplasmids, pQBR57 and pQBR103.

**Figure 1.**
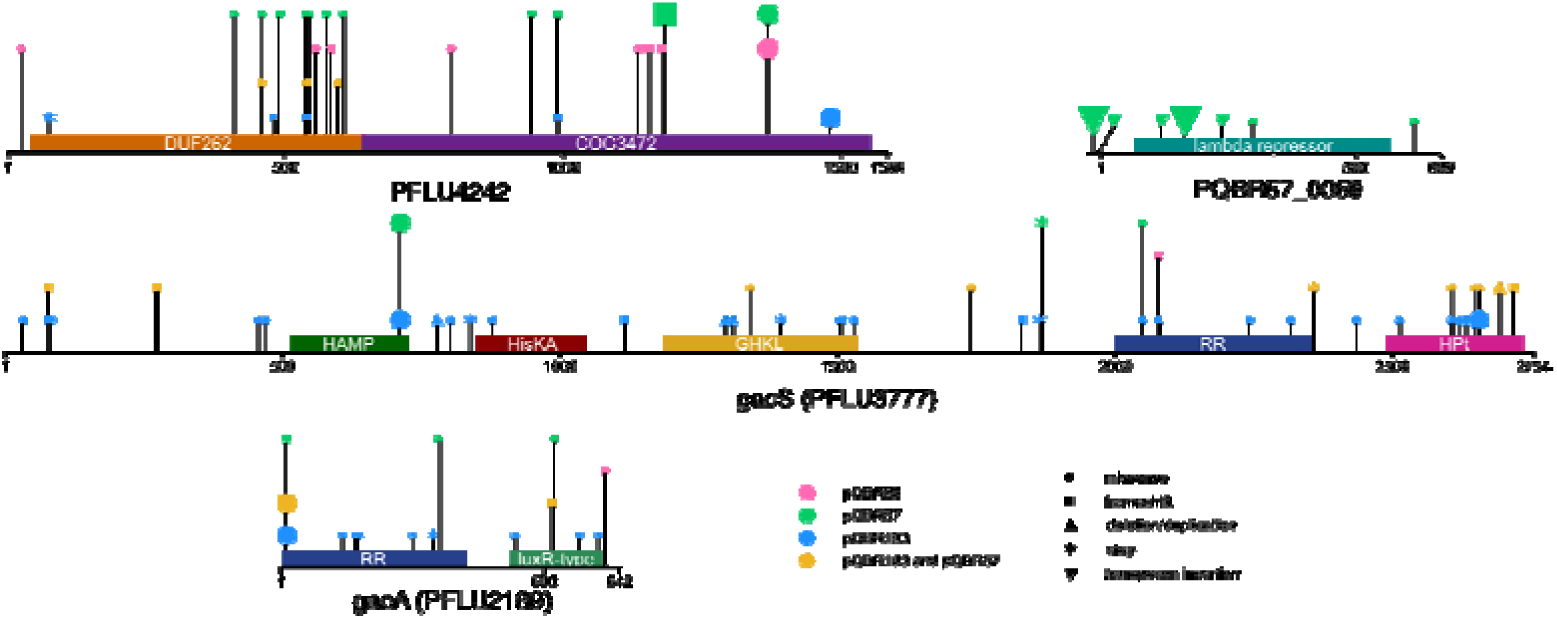
Targets of compensatory mutation. Diagrams show the four genes associated with parallel compensatory mutations from Harrison et al. [33]; Hall et al. [29,39,52] and Carrilero et al. [37]. Symbols indicate the location of the mutation in the nucleic acid sequence. The colour of the symbol indicates the plasmid with which that mutation was associated, and the shape of the symbol indicates the type of mutation. Amino acid-identical mutations (or identical insertion sites) that emerged in parallel in >1 population for that plasmid are shown as a larger symbol. The full list of mutations across these studies, as well as a discussion of how these genes were identified, is provided in **Supplementary Information.**

### Mutations to PFLU4242, to gacS, or to pQBR57_0059 ameliorate the costs of plasmid carriage

First, to test whether *PFLU4242* and *gacS* mutations indeed had a general effect on plasmid cost, we generated pQBR57 and pQBR103 transconjugants in either wild-type bacteria or in mutant backgrounds where we had deleted *PFLU4242* or *gacS*, and measured fitness by direct competition against a plasmid-free isogenic strain (**Fig. 2A**). Knockouts were used since the diversity of naturally-arising mutations to each target (including deletions) was indicative of loss-of-function (**Fig. 1**). Consistent with our hypothesis, we detected a significant effect of mutation on the fitness costs of plasmid carriage (linear mixed-effects model (LMM), plasmid:host interaction likelihood ratio test (LRT) *χ*^2^ = 37.3, p < 1e-6). Each plasmid levied a significant cost in the wild-type background, corresponding with previous studies [37, 45]: for pQBR57, 17.8% (*t* = 4.8, p = 0.003); for pQBR103, 52% (*t* = 10.2, p < 0.001). Mutation of *PFLU4242* resulted in substantial amelioration such that we did not detect a significant fitness cost of either plasmid (post-hoc contrasts with multivariate *t* (‘mvt’) adjustment pQBR57 *t* = 0.87, p = 0.99; pQBR103 *t* = 1.19, p = 0.95). Mutation to *gacS* was more equivocal, with more effective compensation of pQBR103 than of pQBR57 [37]. Neither the Δ*gacS* nor the Δ*PFLU4242* mutants had a detectable fitness difference from wild-type (*gacS t* = −0.24, p = 1; *PFLU4242 t* = 0.27, p = 1), indicating that the effects of mutation were due to interaction with the plasmid rather than providing a direct fitness benefit.

**Figure 2.**
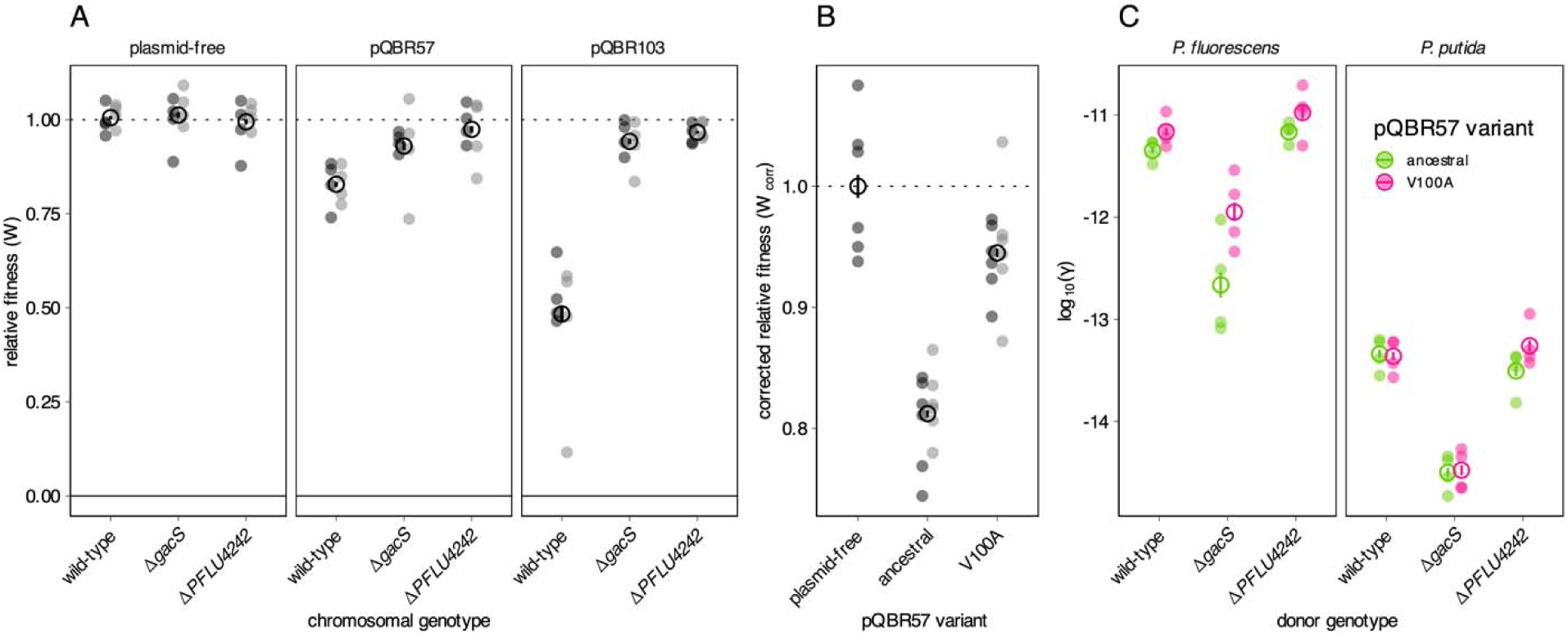
Relative fitness and conjugation rates of plasmid bearing strains and putative compensatory mutations. **(A)** Fitness of wild-type, ΔgacS, and ΔPFLU4242 strains of P. fluorescens SBW25 carrying either no plasmid (left panel), pQBR57 (middle panel) or pQBR103 (right panel). All strains were competed against an isogenic, differentially-marked plasmid-free wild-type P. fluorescens SBW25. Experiments were performed with four independent transconjugant strains as replicates, and were performed twice (light and dark grey). Each measurement is plotted, and the mean across all measurements for each condition indicated with an empty black circle, with error bars corresponding to standard error. **(B)** Fitness of strains with ancestral and V100A variant pQBR57 plasmids. Strains were competed against isogenic, differently-marked plasmid-free strains. Experiments were performed with a total of twelve independent transconjugants into either gentamicin-resistant (dark grey, n=6) or streptomycin-resistant (light grey, n=6) recipients. Relative fitness was corrected for marker effects as competitions were performed with both streptomycin- and gentamicin-resistant test strain backgrounds. For panels **A** and **B**, relative fitness of 1 (horizontal line) indicate no fitness difference between strains. **(C)** Conjugation rates of ancestral and V100A variant pQBR57 plasmids from wild-type, ΔgacS, or ΔPFLU4242 strains of P. fluorescens SBW25 into either isogenic P. fluorescens, or into P. putida KT2440. Experiments were performed with four independent transconjugant strains as replicates. Conjugation was measured according to Simonsen et al. [55]. For all panels, each measurement is plotted, and the mean across all measurements for each condition indicated with an empty circle, with error bars corresponding to standard error.

To investigate the role of PQBR57_0059, we focused our attention on two sequenced, evolved pQBR57 variants clones from an evolution experiment [39]. One of these, A.01.65.G.002 (‘anc’), was identical to the ancestor, while the other B.09.65.G.029 (‘V100A’) had acquired only a single basepair mutation across its 307,330 bp length, resulting in a Val100Ala mutation in the predicted C-terminal domain of PQBR57_0059. By conjugating the A.01.65.G.002 (‘anc’) and the B.09.65.G.029 (‘V100A’) evolved pQBR57 variants into the ancestral strain of *P. fluorescens* SBW25, and each of the chromosomal mutants, we could specifically measure the effects of this mutation (**Fig. 1B**). Competitions showed that the V100A mutation largely ameliorated plasmid cost, (ANOVA, effect of plasmid F_2,27_ = 49.8, p = 8.65e-10; post-hoc contrasts with mvt adjustment anc vs. V100A, p < 1e-8; plasmid-free vs. V100A, p = 0.037), but that possessing this plasmid mutation provided no additional benefit in either the *gacS* or the *PFLU4242* mutant backgrounds (posthoc contrasts, effect of plasmid mutation given chromosomal mutation (|*t*| < 1.7, p > 0.74) (**Fig. S1**). We also tested 10 other naturally-emerging PQBR57_0059 variants and likewise found that, in general, plasmids with a disrupted PQBR57_0059 allele were significantly better at persisting (LMM, effect of PQBR57_0059 disruption *χ*^2^ = 5.6, p = 0.02, **Fig. S2**, **Supplementary Text**), though variance was high between evolved plasmids, probably due to second-site mutations in pQBR57.

Together, these data confirm that mutation to the chromosomal gene *PFLU4242* effectively ameliorates the cost of both pQBR57 and pQBR103 (as we previously showed both for these plasmids [37] and for pQBR55 [29]) and similarly, that mutation to the plasmid gene *PQBR57_0059* ameliorates the cost of pQBR57. As megaplasmid cost can be ameliorated by disruption to just one gene (*PFLU4242*, *gacS*, or *PQBR57_0059*) our results suggest that the major burden of these megaplasmids comes not from the maintenance of plasmid DNA but rather from an interaction that depends on the functioning of specific genes.

### Plasmid compensatory evolution need not negatively affect plasmid transmission

Next, we tested whether the different modes of plasmid compensatory evolution affected conjugation rates. For this, we focussed on pQBR57, since both chromosomal and plasmid-borne genes are implicated. We measured both intraspecific (between isogenic *P. fluorescens* SBW25 strains) and interspecific (to *P. putida* KT2440) conjugation (**Fig. 1C**). Consistent with previous measurements made in soil [19], conjugation into *P. putida* occurred much less readily than between *P. fluorescens* (linear model, effect of recipient F_1,43_ = 925, p < 1e-8. The Δ*gacS* mutation had a significant effect, reducing both intraspecific and interspecific conjugation efficiency by approximately 10-fold (effect of chromosomal mutation F_2,43_ = 110, p < 1e-8). However, neither the Δ*PFLU4242* nor the PQBR57_0059_V100A mutations significantly reduced conjugation, within or between species (p > 0.14 for both modes of compensation). This suggests that there is no necessary trade-off between vertical and horizontal modes of plasmid replication, and that compensatory mutations can potentially enhance both maintenance and spread of conjugative plasmids.

Interestingly, our previous study [33] that identifed GacA/S mutation as a route to pQBR103 amelioration did not find a significant reduction in conjugation rates in the evolved lineages. However, the lack of significance in that study was driven principally by replicates that had mutations in PFLU4242 rather than GacA/S, consistent with our findings with the clean knockout strains reported here.

### Distinct pQBR plasmids have common and divergent effects on chromosomal gene expression

The observation that mutations to *PFLU4242* or *gacA/S* ameliorate the costs of different pQBR plasmids suggests that there are convergent physiological responses to the acquisition and compensation of these different plasmids. To understand these responses and their resolution, we performed RNAseq on plasmid-free bacteria, and from bacteria carrying either pQBR57 or pQBR103, either without amelioration mutations, or with the Δ*PFLU4242* or Δ*gacS* knockouts, or the PQBR57_0059 V100A mutation.

First, we analysed the effect of plasmid acqusition without amelioration. The different plasmids had some common and some divergent effects on chromosomal gene expression (**Fig. 3**, Fig. S3). pQBR57 affected the expression of 398 genes (FDR<0.05), with 47 upregulated >2x and 60 downregulated >2x. pQBR103 affected expression of 254 genes, with 77 upregulated >2x and only 11 downregulated >2x. GO terms enriched amongst the upregulated genes for both plasmids included those associated with the SOS response and signal transduction. However there were divergent effects of each plasmid, particularly for 33 genes which were upregulated by pQBR103 but downregulated by pQBR57. Carbohydrate metabolism was overrepresented in this set (7/33 GO:0005975 “carbohydrate metabolic process,” p_adj_ = 0.008) suggesting that while there was a common response to megaplasmid acquisition, each plasmid also had specific effects.

**Figure 3.**
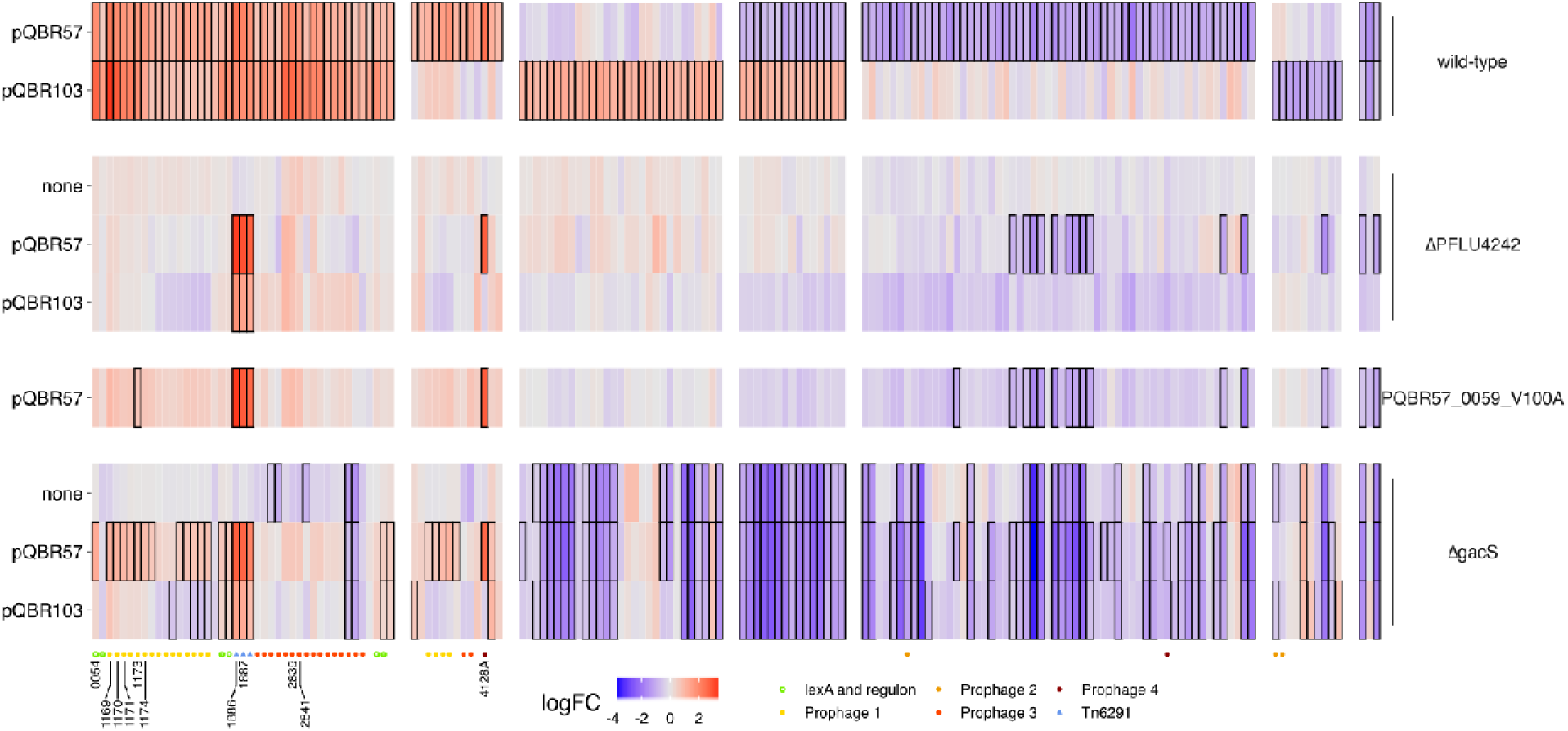
Common and divergent effects on chromosomal gene expression following plasmid acquisition and amelioration. Each column in the heatmap refers to a differentially-expressed (FDR<0.05, absolute log_2_ fold-change > 1) gene following acquisition of either plasmid. Note: there were no additional genes meeting these criteria in the ΔPFLU4242 or PQBR57_0059_V100A ameliorated strains, however, there were 401 additional differentially-expressed genes as a result of gacS deletion, which have not been shown for clarity. Columns have been grouped by pattern of response to pQBR57 and pQBR103. A coloured symbol below each gene indicates whether it is associated with a chromosomal MGE and/or the SOS response. Red indicates increase and blue indicates decreases in expression; cells with a solid border had FDR<0.05 for that specific condition. Genes selected for cloning are labelled.

A closer examination of the differentially-expressed genes shows a set of 50 genes that were at least two-fold upregulated by both plasmids. Many of these were co-localised in the genome, indicating co-regulation, and the possible activation of particular operons by pQBR103 and pQBR57. Of these 50 genes, 31 were in putative prophages, while 3 were in the Tn6291 transposon [39]. Since prophage induction is a signature of the SOS response, and the *P. fluorescens* SBW25 *lexA* homologues *PFLU1560* and *PFLU3605* were also upregulated by both pQBR57 and pQBR103, we investigated how many of the 16 remaining upregulated genes were located downstream of a predicted LexA binding site. By scanning the genome for the conserved pattern CTGKMTNNWHDHHCAG [56] we identified 12 genes <151 bp dowstream of a predicted LexA site, of which 5 were upregulated by both plasmids. The common transcriptional response to pQBR57 and pQBR103 acquisition by *P. fluorescens* SBW25 can thus be characterised as activation of the SOS response and chromosomal mobile genetic elements, which constitute 42/50 (84%) of the >2-fold upregulated genes. This is not the same as the transient SOS response caused by plasmid acquisition reported by Baharoglu et al. [57], as the strains used in these experiments are many generations removed from the original transconjugant due to periods of growth during colony isolation and culture.

Plasmid genes were transcriptionally active. pQBR103 carries 478 predicted ORFs, while pQBR57 carries 426; for both plasmids the majority of ORFs have unknown function. In the ancestral host, expression of pQBR103 ORFs ranged from 0.9–2323 transcripts per million (mean 87, median 39), while expression from pQBR57 ranged from 1.8–2532 (mean 132, median 43). In general, plasmid-borne genes were expressed at levels approximately 2x higher than those of the chromosome, after accounting for gene length (**Fig. 4**). In both cases, the most highly-expressed genes from the plasmids were uncharacterised proteins (**Fig. S4**).

**Figure 4.**
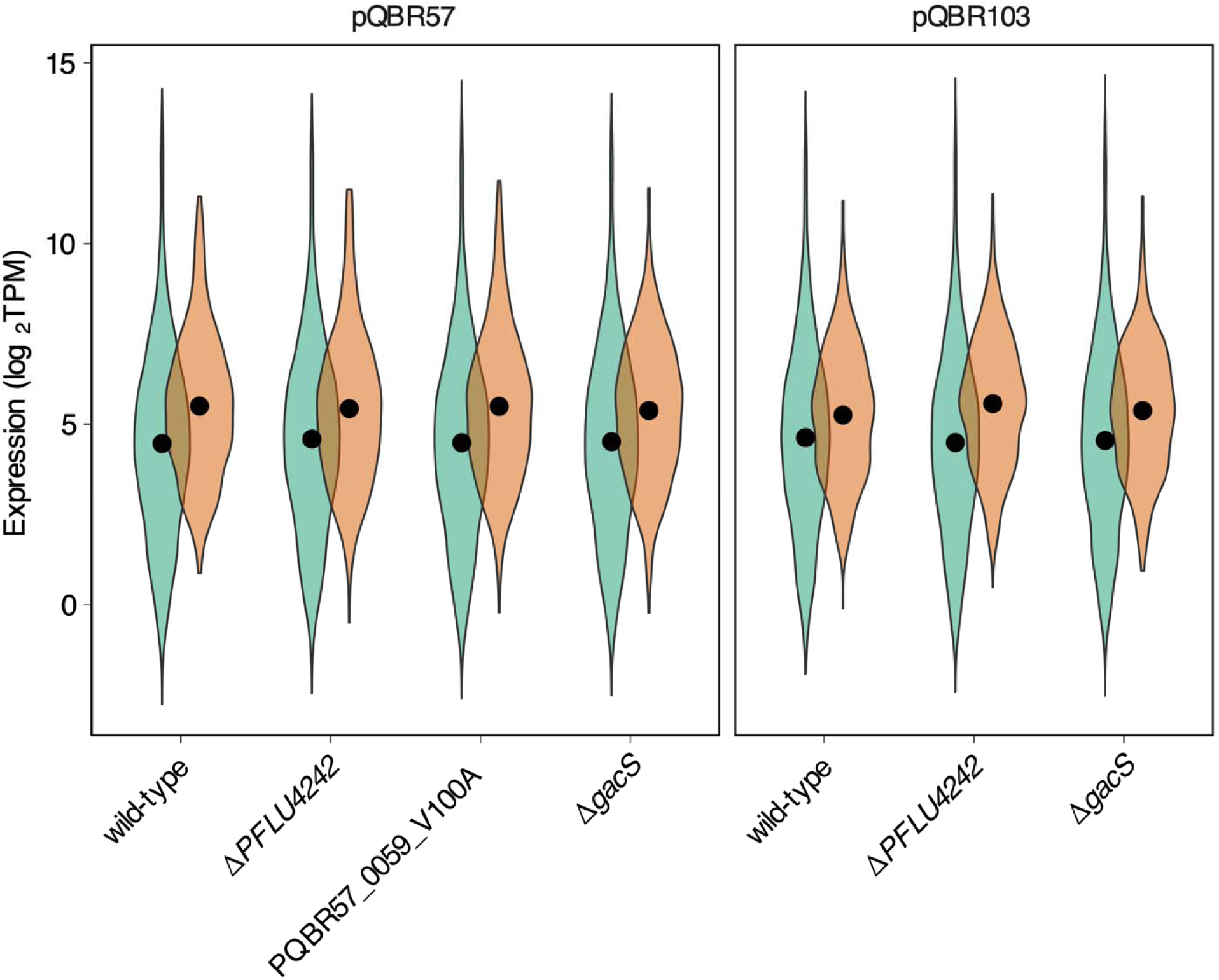
Megaplasmid genes are expressed, and overall levels of expression remain the same regardless of amelioration. Violin plots show log-transformed transcripts-per-million (TPM) of chromosomal (green) and plasmid (orange) genes, averaged across experimental replicates. Solid dots indicate the grand mean for that replicon and condition, calculated from log-transformed values.

**Figure 4.**
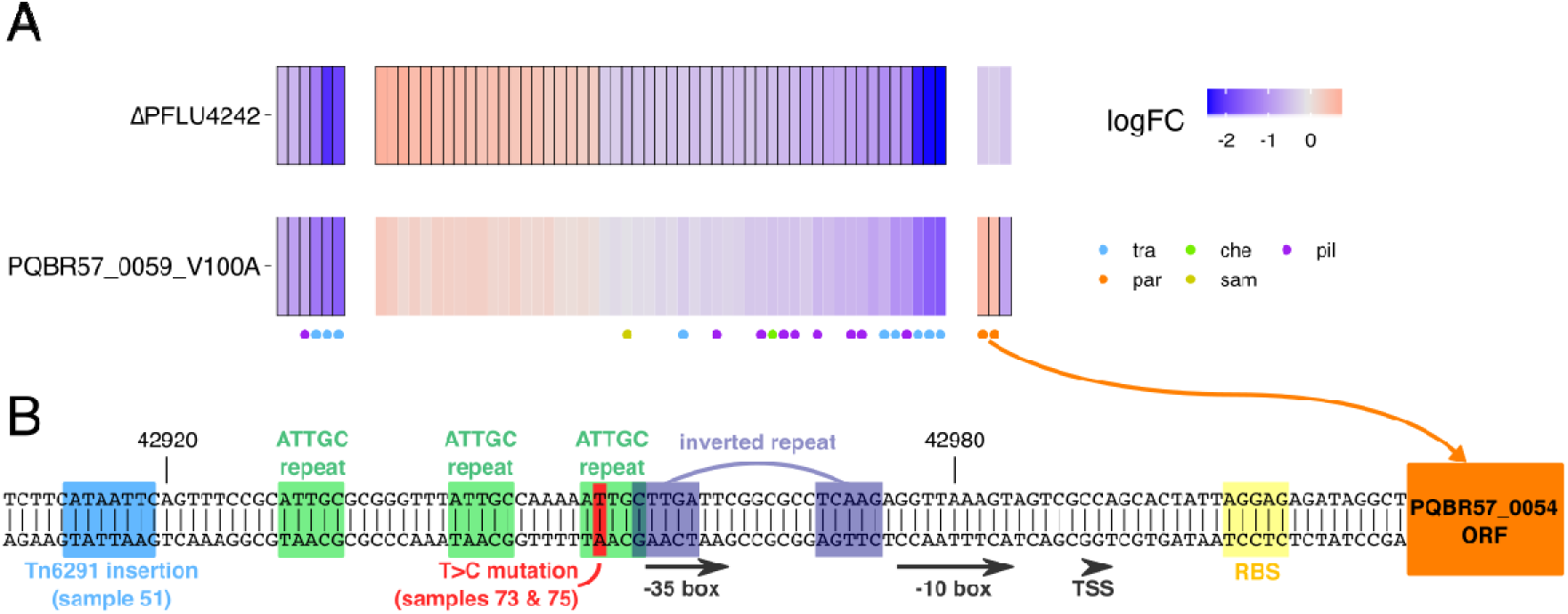
Common and divergent effects on pQBR57 gene expression following different modes of compensatory evolution. **(A)** Heatmap of differentially-expressed (FDR<0.05) genes on pQBR57 following either chromosomally-located amelioration via ΔPFLU4242, or plasmid-located amelioration via the V100A mutation to PQBR57_0059. A coloured symbol below each gene indicates the predicted plasmid region affected. Red indicates increase and blue indicates decreases in expression; cells with a solid border had FDR<0.05 for that specific condition. **(B)** The region upstream of the predicted pQBR57 par genes. Predicted transcription and translation factor binding sites are highlighted, as are identified repeats, and mutations identified by Hall et al. [39]. TSS = Transcription Start Site; RBS = Ribosome Binding Site.

### Plasmid amelioration has broadly similar effects on transcription regardless of mechanism

We then investigated how compensatory mutations affected the changes in transcription caused by plasmid acquisition.

### Effects of PFLU4242 on plasmid cost and amelioration

Without pQBR57 or pQBR103, disruption of *PFLU4242* had no detectable effect on chromosomal gene expression compared with wild-type bacteria (p_adj_ > 0.05 for all genes). However this same mutation in plasmid-bearing bacteria had a striking effect on plasmid-induced changes in gene expression: the reversion to ancestral levels of almost all (pQBR57: 376/398, 94%; pQBR103: 251/254, 99%) of the chromosomal genes differentially regulated following plasmid acquisition (**Fig. 3**), including all of the SOS-associated genes upregulated by both plasmids, and the majority of plasmid-specific differentially-expressed chromosomal genes.

Plasmid genes were also differentially expressed following amelioration by PFLU4242 mutation, but only a subset of them (pQBR57: 57/426; pQBR103: 85/478). The responses of pQBR57 and pQBR103 were largely divergent, with PFLU4242 disruption generally resulting in downregulation of pQBR57 genes, particularly those encoding putative conjugation apparatus (*tra*) and a type IV pilus (*pil*) (**Fig. 5**), while pQBR103 genes were upregulated by PFLU4242 disruption (including *tra* and *pil*) (**Fig. S6**). Though modulation of plasmid gene expression is likely to have some effect on cell physiology, these divergent responses lead us to conclude that the effects of plasmid transcriptional burden or plasmid gene expression *per se* are not primarily responsible for the costs of these plasmids, since the phenotypic costs of carriage could be abolished by *PFLU4242* disruption without major convergent effects on plasmid gene expression.

**Figure 5.**
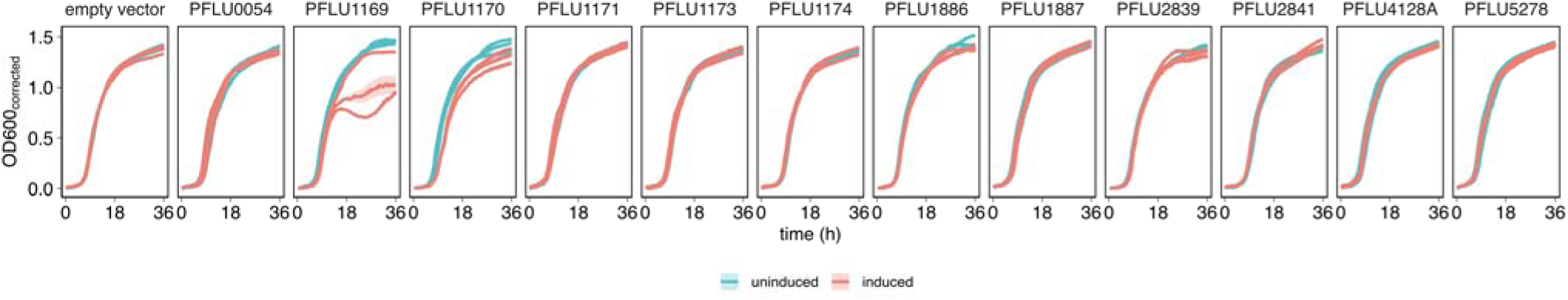
Expression of genes upregulated by megaplasmid plasmid acquisition imposes a fitness cost. Different panels indicate cloned genes. Each vector was transformed into P. fluorescens three times independently, and the growth of each transformant was tested with and without IPTG induction four times, with the mean (solid line) and standard error (shaded area) calculated and plotted for each transformant separately, under 100 µg/ml IPTG (red) or uninduced (blue) conditions. One of the three pME6032::PFLU1169 transformants did not show a large growth defect, but resequencing this plasmid from P. fluorescens revealed a 16-bp deletion in the insert in this replicate. No other mutations were identified in the other replicates, nor in any of the other inserts.

Notably, expression of the three genes from Tn6291 was unaffected by PFLU4242 mutation amelioration for either plasmid. These genes (*PFLU1886-1888*) are hypothesised to encode the transposase subunits for Tn6291. The fact that these genes remain upregulated in the compensated strains suggests that these are not causally responsible for the main negative effects of plasmid acquisition, and also suggests that they are activated through a different route to the other plasmid-upregulated genes.

PFLU4242 mutation therefore had the general effect of bringing chromosomal gene expression back to levels similar to plasmid-free bacteria, suggesting wild-type *PFLU4242* is instrumental in interacting with pQBR plasmids to generate a fitness cost.

### Effects of *gacS* on plasmid cost and amelioration

Compared with *PFLU4242*, unpicking the specific transcriptional effects of *gacS* deletion on plasmid carriage was more difficult, as *gacS* deletion by itself caused differential expression of 733 chromosomal genes affecting numerous phenotypes (see also [58]). One hypothesis was that disruption of GacA/S signalling ameliorates the cost of plasmid carriage by independently downregulating genes that were upregulated by the plasmids, thus ‘balancing out’ the effects of plasmid acquisition. However, while we found a number of genes that were differentially expressed in the plasmid bearers and the Δ*gacS* mutant, the direction of exprssion varied between pQBR57 and pQBR103, and of the 50 genes upregulated by both plasmids (logFC>0, FDR<0.05) only 6 were independently downregulated by the Δ*gacS* mutation, providing little support for this hypothesis.

We therefore considered an alternative hypothesis: that there exists an interaction between GacA/S signalling and the pQBR plasmids that is responsible for plasmid cost, and disruption of the GacA/S system prevents this interaction, resulting in amelioration. One pathway for an interaction between GacA/S and the plasmids could be through PFLU4242. The GacA/S system operates by interfering with the activity of inhibitory mRNA-binding Rsm proteins, causing translation. Other studies have identified putative RsmA binding sites in *PFLU4242*, suggesting that *PFLU4242* may form part of the GacA/S regulon [[59]; Jenna Gallie, pers. comm.]. Indeed, we found that in the Δ*gacS* mutant, *PFLU4242* transcription was reduced by approximately 50%, but *gacS* expression was not affected in the Δ*PFLU4242* mutant (**Fig. S5**). These data provide an explanation for some of the convergence between these two mutational routes to compensation, namely, that PFLU4242 is (more) proximally responsible for plasmid cost, and that disruption to GacA/S could reduce plasmid cost by reducing expression of PFLU4242. Consistent with this hypothesis, and with the phenotypic fitness data, GacA/S disruption affected plasmid-induced changes in gene expression by reverting some of the differentially-expressed genes back to ancestral levels, but this effect was incomplete, particularly with pQBR57 (**Fig. 3**). The fact that PFLU4242 is expressed under wild-type plasmid-free conditions also suggests that PFLU4242 by itself is not toxic, and it is interactions with the megaplasmids that underlies the fitness consequences of this gene.

Disruption of GacA/S signalling affected gene expression from both plasmids in a manner similar to PFLU4242 disruption, but there was an additional effect on pQBR57 which showed 26 genes, mainly of unknown function, specifically downregulated in the Δ*gacS* mutant relative to the wild-type (**Figs. S4, S6**). There was no overall pattern in the predicted functions of these genes, other than their location near the putative origin of replication, and start of the sequence, of pQBR57.

The overlap between plasmid and gac-regulated genes likely represents a complex interaction between the plasmids and the gac system. However, the direct effect that GacA/S has on *PFLU4242* transcription can effectively provide a unified explanation for the two chromosomal modes of amelioration, converging on PFLU4242.

### Effects of PQBR57_0059 on plasmid cost and amelioration

Mutations to PQBR57_0059 had a broadly similar effect on chromosomal gene expression to PFLU4242 (**Fig. 3**). Each amelioration had similar effects on gene expression from pQBR57 (**Fig. 5**), though the effects of the chromosomal mutation were clearer, with many genes trending in a similar direction with the plasmid-borne mutation but not exceeding the FDR<0.05 significance threshold. One clear exception were the *par* genes, which were upregulated by the PQBR57_0059_V100A mutation but not by the ΔPFLU4242 mutation (see below).

### A mechanism for plasmid cost and its amelioration

Together, our transcriptomic data indicates a set of genes that are upregulated by both plasmids, and then downregulated by the different pathways of compensatory evolution. Expression of these genes therefore correlates with the phenotypic fitness costs of plasmid carriage. These genes may be responsible for physiological disruption and fitness costs, but it is also possible that their expression is a downstream effect of other disruptive interactions emerging as a consequence of plasmid acquisition. To test whether expression of these genes causally affects bacterial fitness, we selected 12 candidates for further investigation. Eight candidates were selected as genes which were >2x upregulated by both plasmids in the wild-type but not the Δ*PFLU4242* background, and four additional genes that did not meet these criteria were also selected as controls (**Fig. 3**). The ORF of each gene was cloned into the inducible *Pseudomonas* expression vector pME6032 [60], and the effects of expression on growth were tested by comparing growth curves between induced and uninduced cultures (**Fig. 5**).

We identified one gene, *PFLU1169*, which, when expressed, resulted in a considerable reduction in growth representative of a large fitness cost (LMM effect of insert *χ*^2^ = 49.2, p < 0.0001; Dunnet’s post-hoc test *PFLU1169* vs. empty vector t_26_ = 5.36, p < 0.0001). Indeed, this gene was difficult to clone and express as growth in culture imposed strong selection for mutation of the insert or loss of the expression vector. *PFLU1169* is part of the *P. fluorescens* SBW25 ‘prophage 1’ locus, however, further investigation revealed that this prophage is in fact a tailocin: a protein complex related to a phage tail which acts as a bacteriocin against competitors [61]. *PFLU1169* encodes the holin, expression of which, in concert with other prophage genes, causes cell lysis and tailocin release [62]. The prophage 1 locus is itself regulated through the SOS response, as a LexA binding site was found upstream of this operon. *PFLU1170*, another ‘prophage 1’ gene, had a more marginal effect (*PFLU1170* vs. empty vector t_26_ = 3.2, p < 0.03). Overexpression of any of the remaining 10 genes had no detectable effect (p > 0.99 for all comparisons) under the tested conditions.

We have therefore identified one causal link between plasmid-induced gene expression changes and fitness costs. Plasmid acquisition activates the SOS response, which activates the ‘prophage 1’ tailocin, causing increased cell permeability and lysis. Lysis is likely to be stochastic, the result of which is an apparent reduction in fitness at the population level.

### pQBR57_0059 interacts with other plasmid genes to generate a fitness cost

PQBR57_0059 is predicted to encode a DNA-binding protein of the lambda-repressor family (HHPRED top hit 3BDN ‘lambda repressor,’ p = 4E-23, [63]). We therefore considered it unlikely that PQBR57_0059 was a toxin that directly interfered with cellular physiology. Instead, we predicted that it affected the expression of other genes, either on pQBR57, on the chromosome, or both, resulting in a negative effect on competitive fitness. To distinguish between these alternatives, we cloned either the ancestral-type (‘anc’) or mutant (‘V100A’) variants of the PQBR57_0059 open reading frame (ORF) under the control of the P_lac_ promoter, which is constitutively active in *Pseudomonas*, on the expression vector pUCP18 [41] and introduced these variants to *P. fluorescens* SBW25. We then introduced either the evolved variant of pQBR57 with the V100A mutation, or the unmutated variant. If the costs associated with wild-type PQBR57_0059 were due to direct interaction with the cell, or solely through effects on chromosomal gene expression, fitness costs would be apparent by expressing the ancestral type variant from pUCP18. However if other megaplasmid genes were involved, then pQBR57 would be required to recapitulate the effect, and the wild-type *PQBR57_0059* would complement the otherwise costless V100A pQBR57 variant.

Expressing just the wild-type PQBR57_0059 from pUCP18 was insufficient to generate a fitness cost (**Fig. 6**; LMM, post-hoc tests for pUCP18 variants without pQBR57 p > 0.91 for all comparisons), and other genes on pQBR57 were required for PQBR57_0059 to be costly (interaction between pUCP18 and pQBR57 variant x = 29.4, p = 1e-4). In fact, expression of PQBR57_0059 from pUCP18 appeared to decrease the fitness of pQBR57-bearers even further (effect of anc vs. no insert p = 0.0078), probably due to the increased effect of higher PQBR57_0059 expression from the multi-copy expression vector. Therefore, the fitness effects of PQBR57_0059 are likely to emerge from its effects on other plasmid genes, hinting that a more complex interaction between the plasmid and chromosome is responsible for the cost of pQBR57.

**Figure 6.**
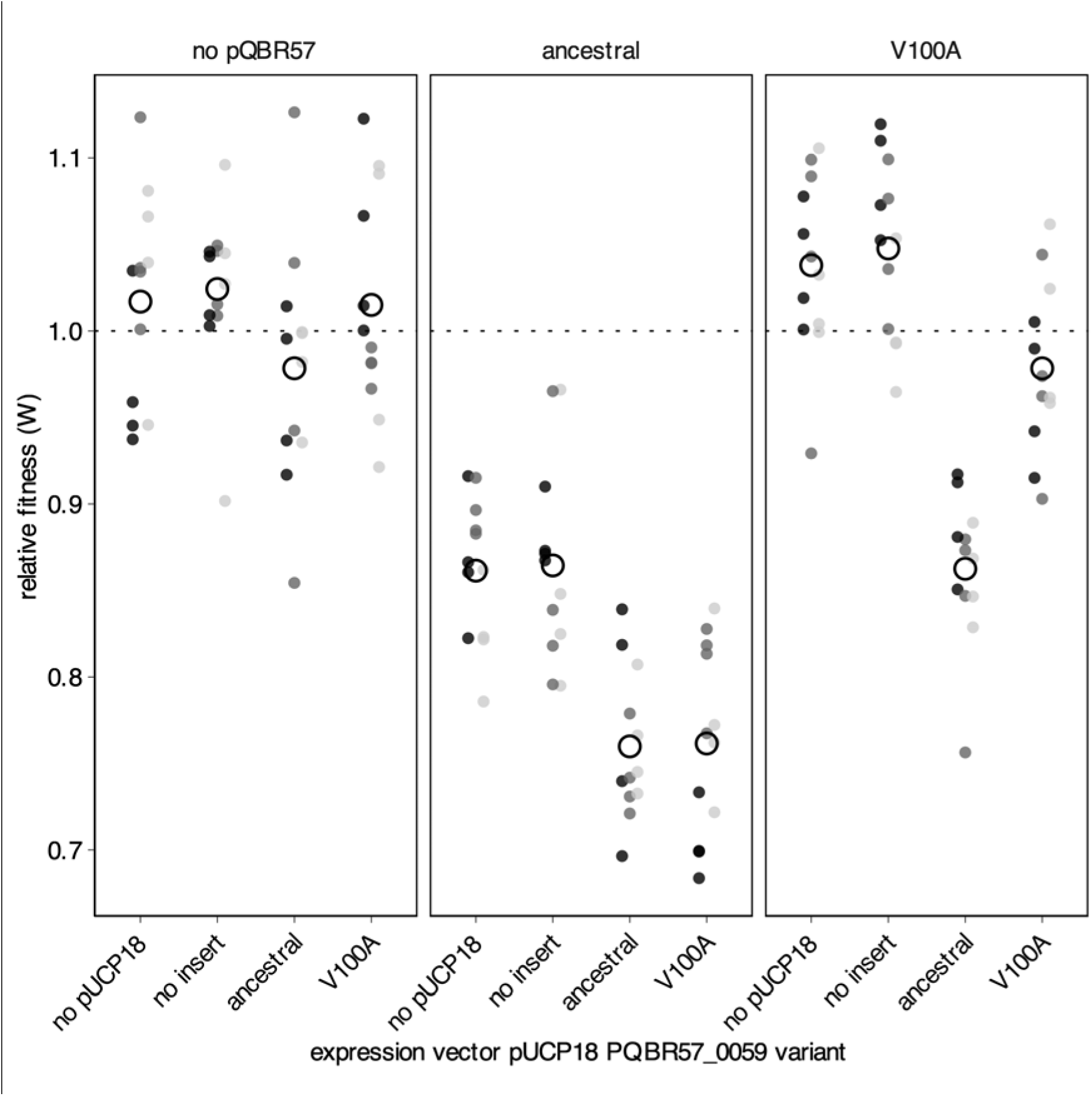
Relative fitness as determined by direct competition for P. fluorescens SBW25 overexpressing either the ancestral or the V100A variant of PQBR57_0059 (or no-insert expression vector, or no-vector controls), and carrying either no pQBR57 or the ancestral or V100A variants of pQBR57. Competitions were performed with three independent transformants/transconjugants indicated by shades of grey, and the fitness of each transconjugant/transformant was tested four times. For the ‘none/no-pQBR57’ control treatment, all measurements were taken on the same strain.

We therefore investigated plasmid-borne genes that were specifically differentially-expressed by the PQBR57_0059 amelioration. Amongst the plasmid genes differentially expressed by amelioration, only three were specifically affected by pQBR57_0059 (**Fig. 4A**). One, PQBR57_0215, was down regulated in a similar direction by the Δ*PFLU4242* mutation (0.05 < p_adj_ <0.1) and thus is likely to be a general response to amelioration. However the other two, which form an operon PQBR57_0054-0055, were exclusively upregulated by pQBR57_0059 disruption. These genes are predicted to encode a ParAB system responsible for efficient segregation of plasmids to daughter cells (HHPRED top hit for PQBR57_0055 6SDK ‘Spo0J/ParB’ p = 2E-24; PQBR57_0054 matches 2OZE_A ‘ParA/Walker-type ATPase’ p = 4.2E-25). Examination of the region upstream of this operon reveals three ATTGC repeats and an inverted repeat (CTTGA(N)_9_TCAAG) that might interact with pQBR57_0059, which is predicted to bind DNA (**Fig. 4B**). Interestingly this region was also the target of parallel mutations in a previous evolution experiment [39]: in one population, evolved pQBR57 carried by *P. putida* had acquired an insertion of Tn6291 transposon 78 bases upstream from the predicted transcriptional start site, while in another replicate, evolved plasmids carried both by *P. fluorescens* and *P. putida* gained a single T>C transition 38 bases upstream of the transcriptional start site. Together this suggests that pQBR57_0059 acts as a repressor of the *par* locus, and disruption of this repressor, or disruption of its binding site, results in increased transcription. The fact that there is no additive benefit of mutations in PFLU4242 and pQBR57_0059 suggests that increased par gene expression somehow prevents a deleterious interaction with PFLU4242, directly or indirectly, to ameliorate plasmid cost.

### Increased expression of the *par* genes from pQBR57 is sufficient to ameliorate the costs of pQBR57, but not pQBR103

To test the hypothesis that increased *par* gene expression as a consequence of the disruption of the repressor PQBR57_0059 was responsible for ameliorating the costs of pQBR57, we cloned the putative *par* genes PQBR57_0054-0055 into pUCP18 and introduced this vector, or an empty pUCP18 control, into *P. fluorescens* SBW25. We then introduced pQBR57, and measured the effects on relative fitness. Consistent with our hypothesis, overexpression of the *par* genes ameliorated the costs of pQBR57 (**Fig. 7A**; LMM, interaction of pUCP18 insert and pQBR57 carriage *χ*^2^ = 22.2, p < 1e-4; pairwise contrasts no insert pQBR57 vs. *par* pQBR57 *t*_10_ = 5.15, p = 0.004), but had no significant fitness effects by themselves (no insert no pQBR57 vs. *par* no pQBR57 *t*_10_ = 1.82, p = 0.5).

**Figure 7.**
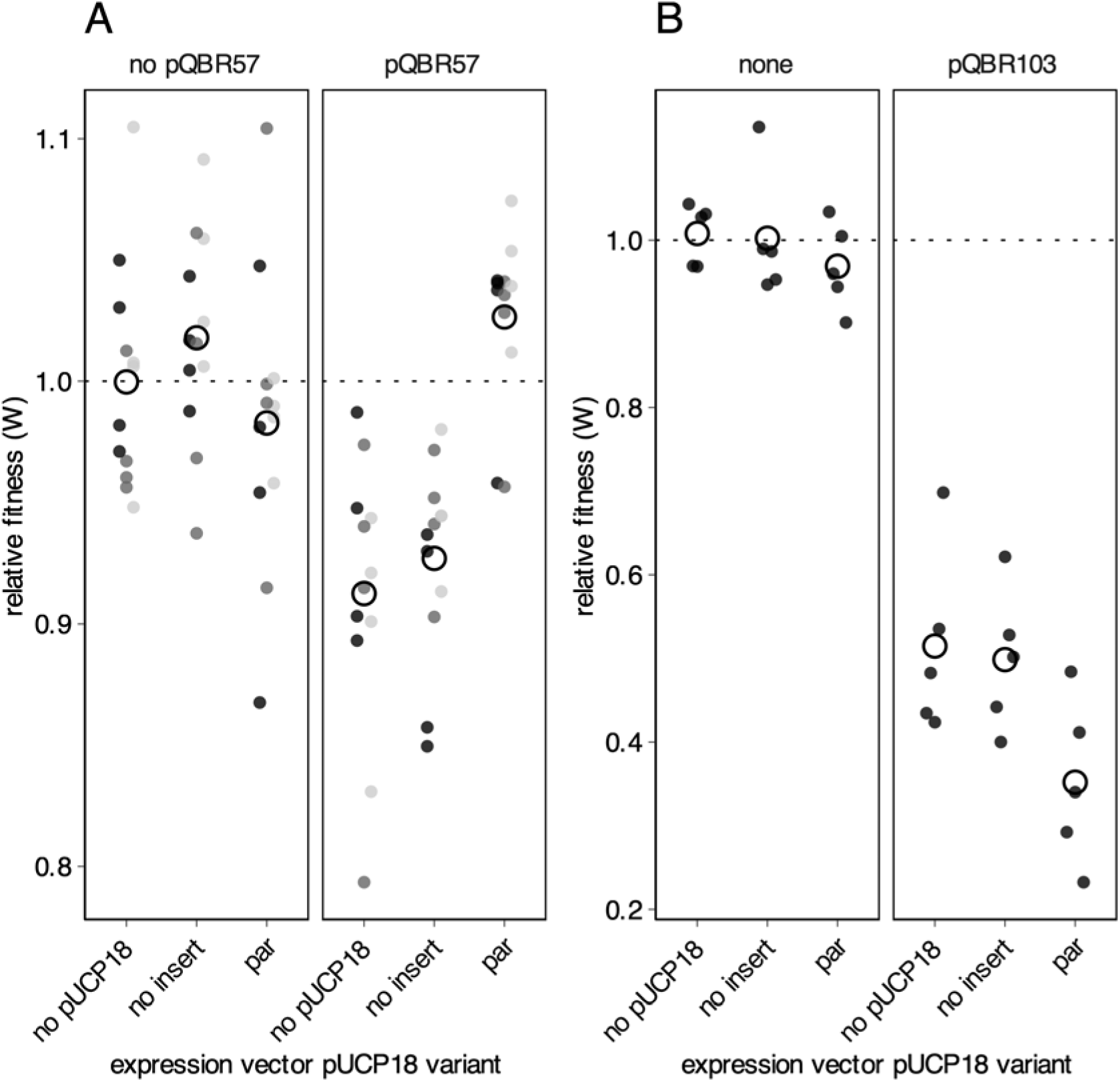
Effects of pQBR57 par gene expession on megaplasmid cost. **(A)** Competitions were performed with three independent transformants/transconjugants indicated by shades of grey, and the fitness of each transconjugant/transformant was tested four times. Empty circles indicate the grand mean for that set of conditions. **(B)** Expression of pQBR57’s par genes in trans exacerbates the fitness costs of the distantly-related megaplasmid pQBR103. Competitions were performed with five independent transconjugants, with each one tested once.

We then investigated whether overexpression of pQBR57’s *par* genes had a general effect on plasmid fitness costs. We introduced pQBR103 to the strains that were overexpressing PQBR57_0054-0055 and again measured relative fitness. In contrast to the amelioration of pQBR57, expression of PQBR57_0054-0055 had no beneficial effect on pQBR103 (LM, pUCP18:pQBR plasmid interaction F_2,24_ = 1.79, p = 0.18; pairwise contrast pQBR103 no insert vs. pQBR103 *par t*_26_ = 2.41, p = 0.12), and we had weak evidence that costs were in fact exacerbated (effect of pUCP18 insert F_2,26_ = 4.4, p = 0.02). Increased expression of pQBR57 *par* therefore reduced the fitness costs of pQBR57, but did not have any ameliorative effect on an heterologous plasmid.

## Discussion

Plasmids are known to impose fitness costs, but the molecular causes of such costs are far from clear. Here we show that, rather than emerging from the general bioenergetic demands of replicating DNA, transcribing into mRNA, or translating into protein [20], the majority of the fitness costs of plasmid acquisition come from specific deleterious interactions between plasmid and chromosomal genes. As a consequence, small mutations — in some cases just single base changes — can be sufficient to ameliorate even a very large plasmid with hundreds of genes. Previous studies have tended to focus on smaller, non-conjugative replicons [25,30,64], or have identified plasmid compensatory mutations with substantial pleiotropic effects: disruption or deletion of conjugative machinery from the plasmid (rendering it immobile) [28, 65]; deletion of large portions of the plasmid [34, 66]; or disruption of extensive and multi-functional regulatory circuits [31,33,67]. In contrast, our data shows that single mutations can enable large, natural, conjugative, plasmids to become rapidly accommodated with few obvious tradeoffs, and furthermore describes the molecular negotiation by which this occurs (**Fig. 8**).

**Figure 8.**
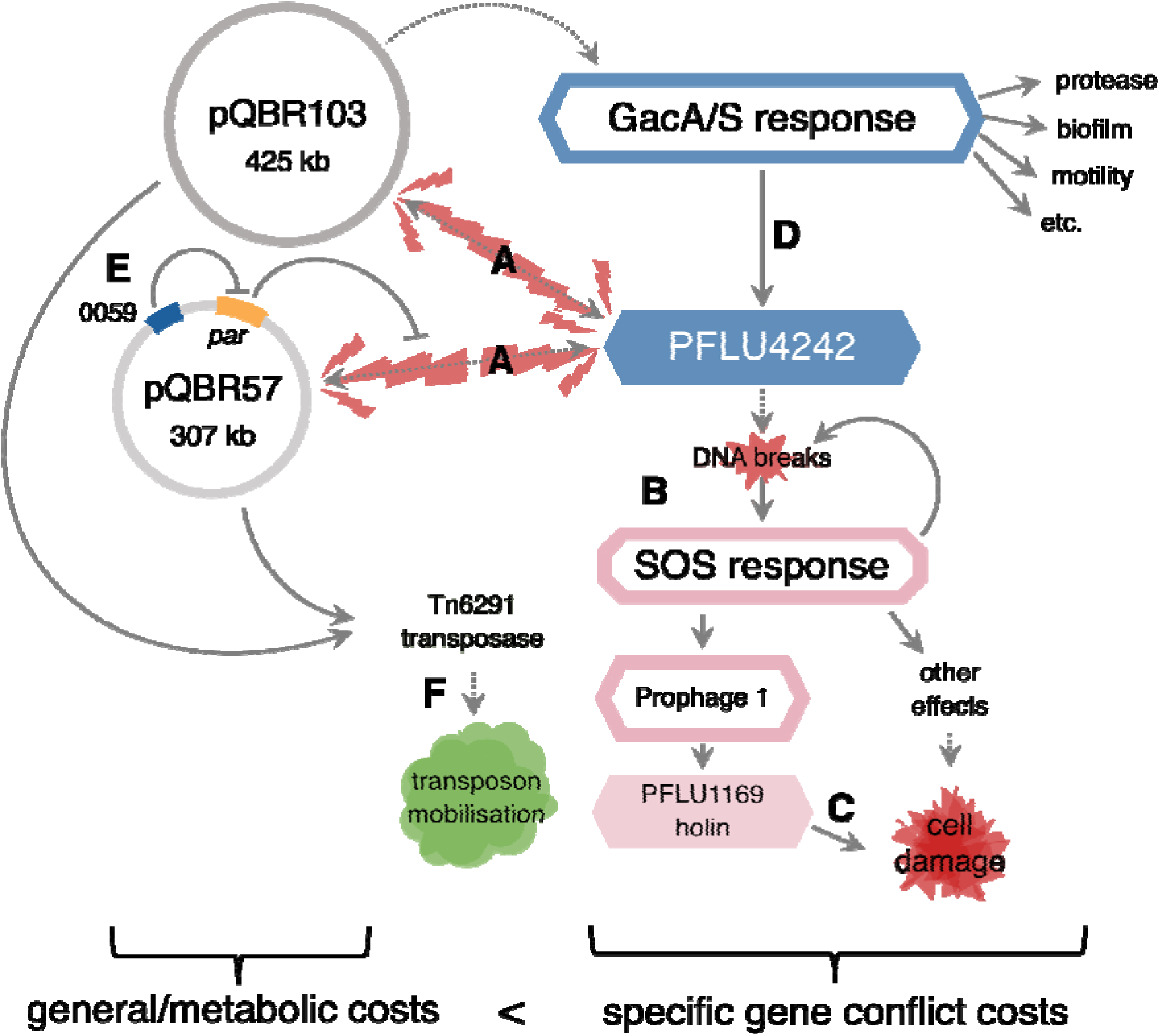
Interactions leading to pQBR plasmid fitness costs. Dotted lines indicate speculative interactions, solid lines indicate experimentally-or bioinformatically-determined interactions (from this study or others). Targets of compensatory mutation are shown in blue. Genes are solid boxes whereas pathways/groups of genes are outlines. **(A)** Both plasmids interact with PFLU4242. **(B)** PFLU4242 activates the SOS response, probably through causing DNA breaks. **(C)** One outcome of the SOS response is expression of PFLU1169 which directly causes cell damage. **(D)** The effects of plasmid interactions with PFLU4242 are exacerbated by GacA/S, which increases expression of PFLU4242. pQBR103 may also interact directly with the GacA/S system. **(E)** Mutation of PQBR57_0059 increases expression of par, which reduces the disruption of pQBR57 but not pQBR103. **(F)**. Both plasmids activate the Tn6291 transpose through a different route to PFLU4242 or GacA/S, which is likely to increase transposition activity and mobilisation of Tn6291. Overall, specific gene conflict costs greatly exceed the metabolic costs of pQBR57 and pQBR103.

Specifically, we have linked the fitness costs of both pQBR57 and pQBR103 to the SOS response in *P. fluorescens* SBW25. SOS response-associated genes comprised 42/50 of the genes upregulated in common, and we found that increasing expression of just one gene in the SOS regulon — the ‘prophage 1’ holin *PFLU1169* — had a severe effect on fitness, implicating the SOS response as a direct proximal cause of plasmid-induced fitness costs. The SOS response has wide-ranging impact on bacterial physiology [68], meaning that *PFLU1169* is unlikely to be the *sole* mechanism of plasmid-induced fitness costs. How is the SOS response triggered by the pQBR megaplasmids? Single-stranded DNA, such as is acquired in the early stages of conjugative transfer, can directly activate the SOS response [57], but as the strains in this study were many generations removed from the original conjugation event due to picking and restreaking of transconjugant colonies, the SOS response in the pQBR/SBW25 system is clearly being perennially activated through a different route. DNA damage is the most likely candidate, implicating the SOS response as essentially an amplifier of disruptions caused by plasmid acquisition. Like an amplification circuit, the SOS response can itself cause DNA fragmentation, forming a positive feedback loop, and further disruption [69].

How does plasmid carriage cause DNA breaks? Clashes between the machineries of transcription and replication put stress on DNA molecules, and such clashes have been shown to drive the fitness costs of the minimal (2.1 kb) plasmid pIND in *E. coli* [64]. In *Shewanella oneidensis*, the replication machinery of the IncP1-beta mini-replicon pMS0506 (13.1 kb) probably triggers the SOS response by binding to the host DNA helicase DnaB and blocking the replication fork, exposing vulnerable ssDNA intermediates to breaks [70]. In *P. aeruginosa* PAO1, the small (5.1 kb) plasmid pNUK73 triggered the SOS response due to increased expression of the plasmid replication machinery, which may cause breaks due to increased stress on DNA molecules within the cell [25]. However, other costly plasmids did not have the same effect on *P. aeruginosa* PAO1, suggesting that SOS response activation was not a general source of plasmid fitness costs in that strain [26]. The fact that different species of *Pseudomonas* had differing SOS responses to the same pCAR1 metabolic plasmid [71] likewise suggests that activation of the SOS response usually occurs in a plasmid-by-host specific manner, again consistent with our hypothesis that plasmid fitness costs emerge principally through specific genetic conflicts.

It is unclear whether the conflicts between transcription and replication seen with small plasmids is responsible for DNA damage for much larger plasmids like pQBR103 and pQBR57. We saw no significant change in megaplasmid copy number [39] or *rep* gene expression when comparing ancestral with ameliorated strains, hinting that other processes may be at work. What is clear is that mutations to a limited number of targets can prevent activation of the SOS response, presumably by preventing DNA breaks from occurring. Principal amongst these mutations, in terms of degree and scope of amelioration, are loss-of-function mutations to PFLU4242. While the molecular and evolutionary functions of PFLU4242 remain unknown, its DUF262 domain pattern resembles that of known endonucleases that target modified phage DNA [72], and related genes are located in known genome defence islands [73]. The hypothesis that PFLU4242 has an nuclease function in phage defence would explain how PFLU4242 disruption might prevent DNA breaks, and also why this gene has been maintained despite having no detectable effect on fitness or gene expression in plasmid-free bacteria in an axenic laboratory setting. Work is ongoing to understand the molecular and evolutionary functions of this gene. The concept of a trade-off between openness to HGT and resistance to phage infection has been explored extensively in the context of CRISPR-Cas and restriction-modification modules [14,74,75]. Our data hints at another mechanism by which genome defence might have side effects: an inappropriately excessive response driving fitness costs, analogous to immune hypersensitivity.

We show that PFLU4242 expression is induced by a global regulator of secondary metabolism, the GacA/S two-component system, and so our data is consistent with the hypothesis that disruption to GacA/S could exert its effect through repression of PFLU4242. It is possible that global regulators have been implicated in plasmid persistance due to their regulatory effects on more proximal causes of plasmid conflict [31, 67]. The combined size of the *gacA* and *gacS* genes (∼3.4 kb) is over twice that of *PFLU4242* (∼1.6 kb) (**Fig. 1**), providing a larger mutational target, which, alongside the mutability of the GacA/S system [76], may explain why GacA/S rather than PFLU4242 was the primary target of mutation in one of our previous studies [33]. However, we have hints that PFLU4242 is not the only mechanism by which pQBR plasmids and the GacA/S system interact. Acqusition of pQBR103 has a similar effect on PFLU4242 expression as deletion of *gacS*, while 26 pQBR57 genes appear to be differentially regulated in the Δ*gacS* background but not in the Δ*PFLU4242* background. Moreover, pQBR103 appears to carry a homologue of the Gac system signal transducer RsmA, in common with pQBR55 but not pQBR57 [45]. Understanding how plasmids may manipulate core pathways in bacterial signalling is an important subject for future investigation [77].

Compensatory mutations occuring on a plasmid are thought to have increased potential to enhance plasmid persistence and accelerate horizontal gene transfer relative to mutations of similar effect on the chromosome, because plasmid-borne compensation is inherited both by daughter cells and by transconjugants [35]. Mutation of the putative lambda-family repressor PQBR57_0059 caused increased expression of the putative partitioning machinery of pQBR57, reducing the cost of that plasmid. The molecular mechanism by which *par* expression reduced pQBR57’s cost remains unclear. Chromosomal ParB is known to bind DNA both specifically (at *parS* sites) and non-specifically, impeding access by DNA-damaging agents like nucleases, forming bridges and loops between distant parts of a DNA sequence, and affecting gene expression [78–83]. The putative ParB of pQBR57 may also act to prevent DNA damage, perhaps by restricting nuclease access to otherwise vulnerable DNA. What, then, is the evolutionary function of PQBR57_0059, in repressing the protective *par* module? Tuning *par* expression for plasmid cost reduction may be a delicate matter, as our data suggests that high levels of pQBR57 *par* expression alone has a cost (as has also been found for chromosomal ParB expression [79, 82]), and may exacerbate the costs of heterologous plasmids as we see for pQBR103. A highly conjugative plasmid such as pQBR57 is expected to frequently find itself in co-existence with other plasmids [84], and PQBR57_0059 may be one mechanism by which pQBR57 can modulate its own activity in multi-replicon cells. Other plasmids are known to carry genes that prevent cellular disruption, for example the anti-SOS genes PsiB [85], or the nucleoid-like proteins H-NS that repress plasmid gene expression [86], reducing costs. In this context, it is interesting that we have witnessed the evolution of this ‘stealth’ function for the pQBR57 *par* genes during the course of our evolution experiment, showing that plasmids can contain the genetic resources necessary to evolve their own amelioration.

The principal group of genes upregulated by both plasmids were located in chromosomal regions annotated as mobile genetic elements. Foremost amongst these were prophages 1 and 3. However, the prophages of SBW25 are not predicted to be intact integrated phage, and rather seem to be relics which have lost the genes involved in DNA replication and packaging [62,87,88]. These prophages are therefore candidate tailocins — phage tails that have been evolutionarily co-opted to act as toxins used in inter-strain competition [61]. To exert their activity, tailocins translated in the cytoplasm must be released by cell lysis [61]. Tailocin production is therefore an example of bacterial spite, whereby an individual suffers a fitness cost to harm a competitor [89]. Spite, like altruism, is evolutionarily stable only within a narrow range of conditions — specifically, where those benefitting from the action are likely to be related, and those targeted by the action are unlikely to be related [90]. Receiving a new plasmid is a clear signal of the presence of an unrelated competitor, and thus an opportune time for spiteful tailocin activation. The SOS response can be considered a strategy against aggressions, rather than a solely defensive reaction [68]. It is therefore tempting to speculate, that, rather than being solely a side effect of DNA damage, expression of tailocins following plasmid acquisition could be an adaptive trait, that both inhibits competitors [90] and potentially also prevents invasion of parasitic plasmids through cell suicide in a manner similar to abortive infection [91].

The other chromosomal mobile genetic element upregulated by both pQBR57 and pQBR103 was the transposon Tn6291. The upregulated genes are thought to encode the transposase, which provide the machinery enabling Tn6291 to relocate both to other chromosomal locations and, importantly, to conjugative plasmids. Tn6291 does not appear to encode conjugative machinery of its own, and therefore depends on other apparatus to move horizontally between bacterial lineages. Indeed, we previously showed using experimental evolution that Tn6291 exploits pQBR57 to move between *Pseudomonas* species in soil [39]. This dependence of Tn6291 on other mobile genetic elements for transmission provides an adaptive explanation for why the transposase genes are sensitive to conjugative plasmid acquisition: the presence of a conjugative plasmid signals the presence of a vehicle for Tn6291 to use. Communication between the pQBR plasmids and Tn6291 appears to be independent of the SOS response, as the Tn6291 transposase remains upregulated in the otherwise ameliorated plasmid-bearing strains. There are numerous adaptive reasons why mobile genetic elements might be expected to communicate with one another, since conflict and collaboration between mobile genetic elements is expected to be a key factor influencing horizontal gene transfer and hence bacterial evolution [e.g. 2,92]. Interpreting this language could prove key to understanding the factors driving the spread of ecologically- and clinically-important traits.

Our work has focused on megaplasmids: large plasmids, generally exceeding 150 kb in size. Megaplasmids are likely to be especially effective vectors of horizontal gene transfer as they can mobilise many traits at once — a particular concern for the spread of antimicrobial resistance because selection with one drug may drive the spread of multi-drug resistance [93]. Similarly, compensation of megaplasmids potentially facilitates the maintenance of many different traits in a lineage, provided that (consistent with our findings here) costs do not emerge from plasmid size *per se*. Advances in long-read sequencing technologies are beginning to expose the diversity, ubiquity, and complexity of megaplasmids in diverse bacterial genera, and therefore our findings are likely to apply beyond this specific case of mercury resistance in a soil Pseudomonad [e.g. 94,95]. For example, megaplasmids pQBR103 and pQBR57 are distant relatives of a large family of clinically important plasmids that appear to be disseminating multi-drug resistance in clinical isolates of *Pseudomonas aeruginosa* from opportunistic infections [96, 97]. In those strains, megaplasmids appear to be stable even without the selective pressure of antimicrobials, suggesting that those strains are either pre-adapted for megaplasmid carriage, or have undergone compensatory evolution.

The fact that plasmid costs emerge principally from specific gene conflicts is not to say that replication and expression of megaplasmid genes *per se* levies no burden, nor that such costs are invisible to selection. Lynch and Marinov [20] calculated the costs of bacterial gene replication, transcription, and translation, and found that the costs of even replicating short pieces of non-beneficial DNA are sufficient for purifying selection in moderately-sized bacterial populations, with the costs of transcribing and translating those genes several orders of magnitude greater. The assays we used to measure fitness have limits, and it is likely that the residual bioenergetic burden of plasmid carriage would be detected by more sensitive methods [98]. Nevertheless, it is clear that such bioenergetic costs are many fold smaller than those that emerge from conflicts between specific genes or their products, and may well be overwhelmed by other selective pressures in natural environments (such as phage [99]) or rendered negligible in the context of horizontal gene transfer [100].

Our findings have important implications for understanding the evolution of bacterial genomes. Fitness costs of plasmid acquisition are, in theory, a barrier to HGT, however, our data suggest that when such costs are caused by specific genetic conflicts they will be readily and rapidly ameliorated by single compensatory mutations thus enabling the long-term maintenance of plasmids in bacterial genomes. This helps to explain why plasmids are so common in bacterial genomes and, moreover, suggests that such compensated plasmid-carrying lineages will become important hubs of HGT within bacterial communities [19, 101]. The fact that plasmid costs depend on specific conflicts is relevant to understanding the dissemination of ecologically and clinically important bacterial traits, such as antimicrobial resistance, because the immediate fitness costs of the encoding plasmids are unlikely to predict their long-term dynamics: purifying selection against plasmids will diminish over time due to rapid compensatory evolution. More generally, the dynamics of mobile elements in bacterial genomes are unlikely to be predictable from their generic properties, and thus more sophisticated analyses of genetic associations in pangenomes [e.g. 102] will be required to predict the patterns and outcomes of gene exchange and how these will shape pangenome dynamics.

## Author contributions

**James P. J. Hall:** Funding acquisition, Conceptualization, Data Curation, Formal Analysis, Investigation, Methodology, Visualization, Writing — Original Draft Preparation, Writing — Review & Editing

**Rosanna C. T. Wright:** Resources, Investigation, Writing — Review & Editing

**Ellie Harrison:** Funding acquisition, Conceptualization, Writing — Review & Editing

**Katie Muddiman:** Resources

**A. Jamie Wood:** Funding acquisition, Conceptualization, Writing — Review & Editing

**Steve Paterson:** Funding acquisition, Supervision, Conceptualization, Writing — Review & Editing

**Michael A. Brockhurst:** Funding acquisition, Conceptualization, Project Administration, Writing — Review & Editing

## Acknowledgements

We would like to thank staff at the NERC Biomolecular Analysis Facility, University of Liverpool, for assistance with the RNA-seq, and Jenna Gallie (Max Planck Institute for Evolutionary Biology) for a list of putative RsmA targets in *P. fluorescens* SBW25.

---

**Supplementary Figure S1.**
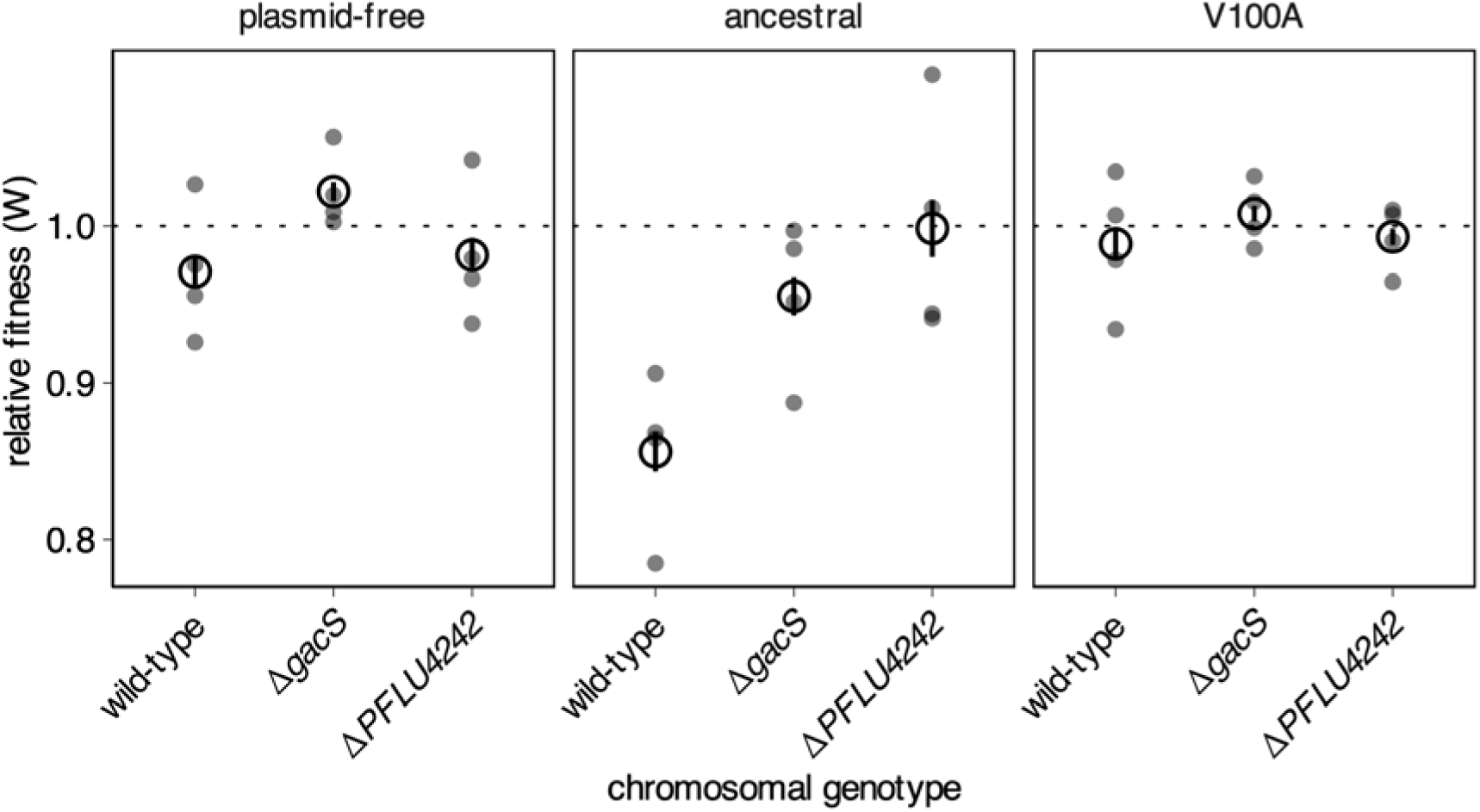
Relative fitness as determined by direct competition for ancestral and V100A variants of pQBR57, in wild-type, ΔgacS, or ΔPFLU4242 strains of P. fluorescens SBW25, against plasmid-free wild-type P. fluorescens SBW25. Competitions were performed with four independent transconjugants. Each measurement is plotted and the mean across all measurements for each condition indicated with an empty black circle, with bars indicating standard error.

**Supplementary Figure S2.**
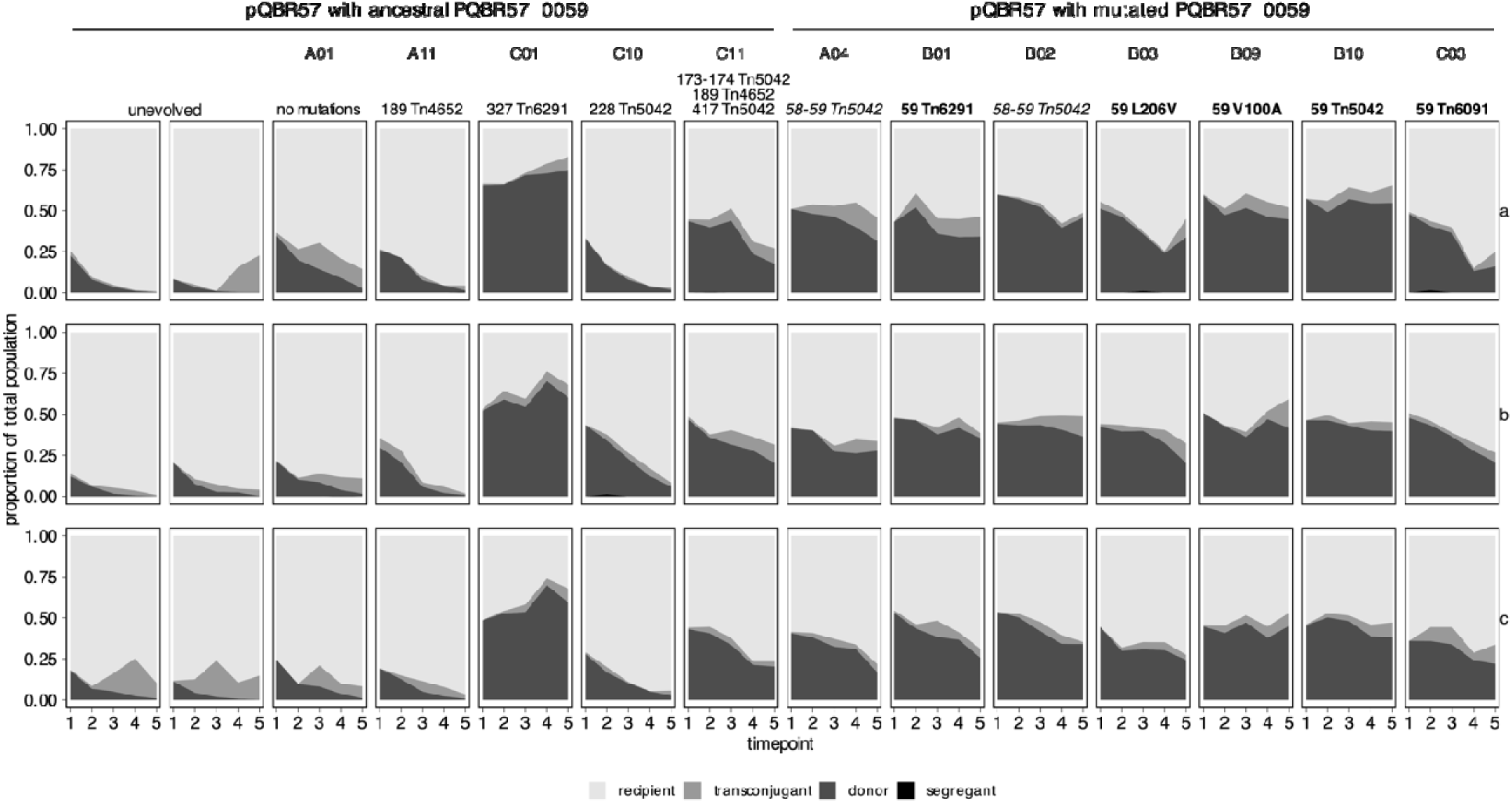
Serial passage of plasmid-bearing strains in competition in soil microcosms. Panels show the relative proportions of different subpopulations over five transfers for three independent replicates (rows a, b, c). Columns correspond to different pQBR57 variants which were conjugated into an ancestral strain for this experiment. Columns are arranged according to whether that variant had a disruption in PQBR57_0059 (right-hand columns) or not (left-hand columns). All detected mutations in that variant are described at the top of the column, with numbers corresponding to the affected locus_tag (with leading zeroes removed for reasons of space). ‘Tn’ indicates a transposon insertion. Mutations in PQBR57_0059 are indicated in **bold**, mutations affecting the region immediately upstream of PQBR57_0059 are in italics. Columns 1 and 2 contain data reproduced from Hall et al. [39].

**Supplemental Figure S3.**
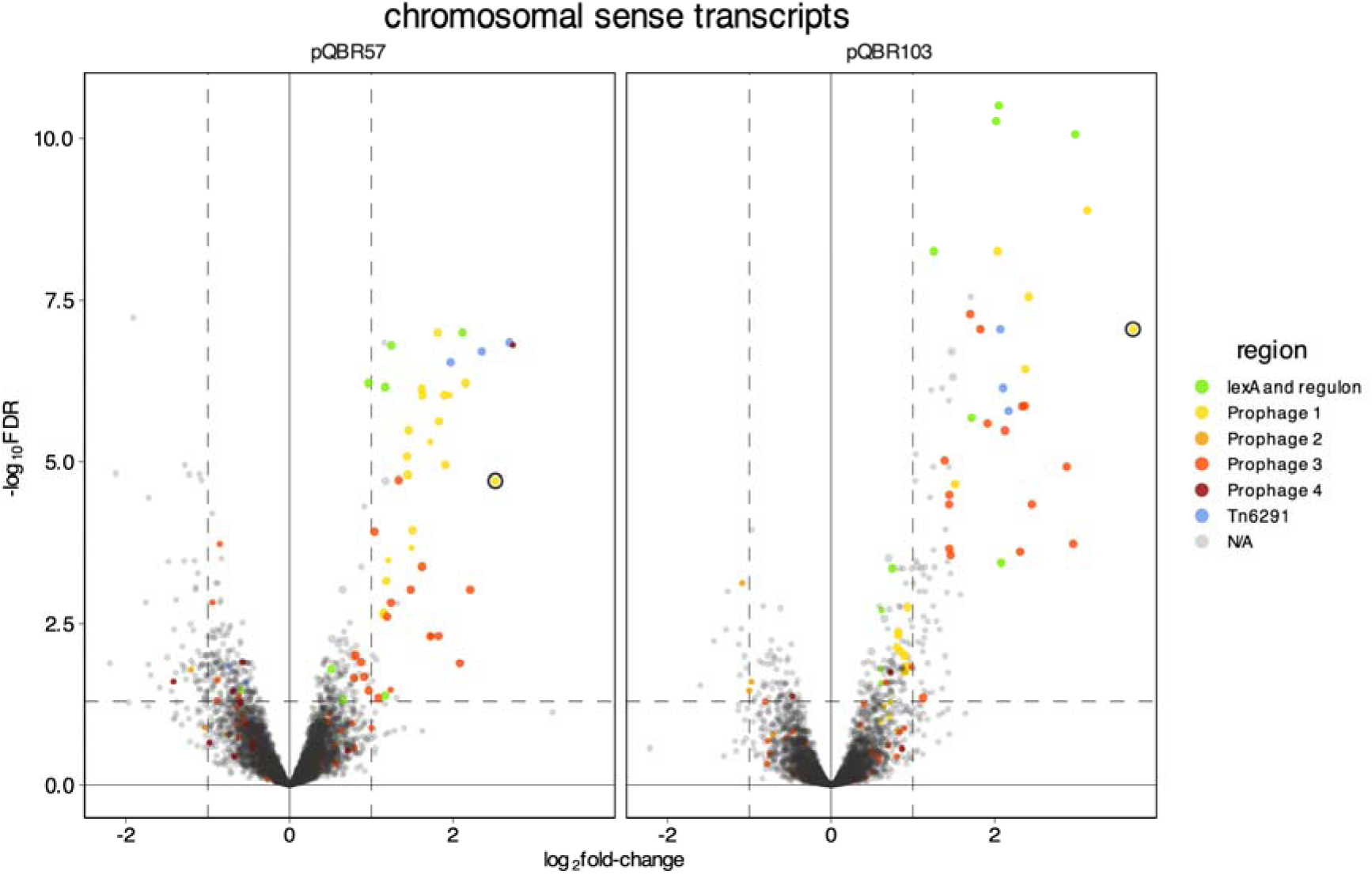
Volcano plots showing changes to gene expression following plasmid acquisition. Points are coloured according to Figure 4. PFLU1169 is highlighted with a black border.

**Supplementary Figure S4.**
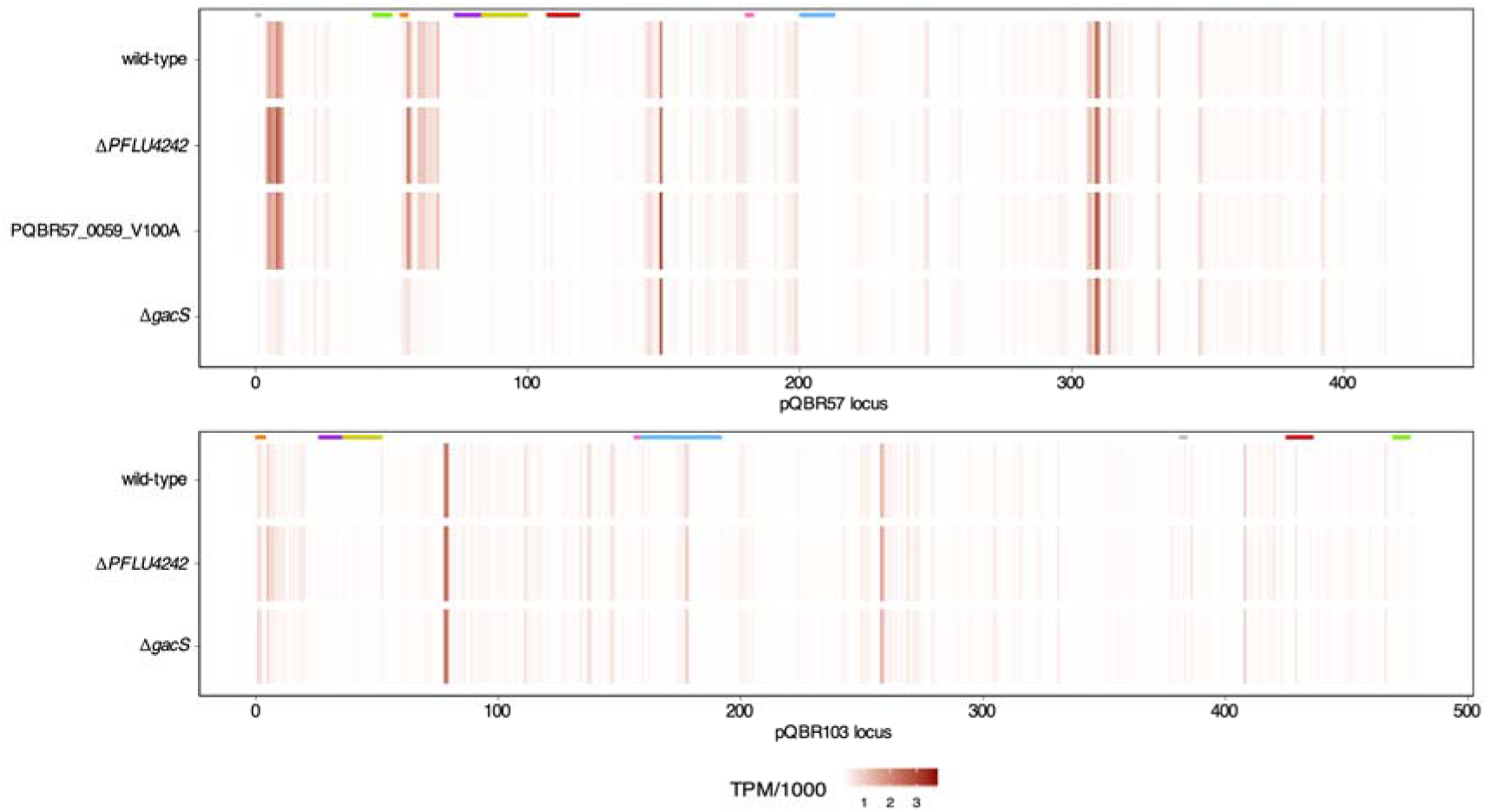
Gene expression varies across the plasmids, with the most highly-expressed genes outwith regions with predicted functions. Heatmaps show level of expression in transcripts-per-million (TPM) across each plasmid under different amelioration conditions. Coloured bars across the top indicate regions annotated in Hall et al. [45]: grey = rep; green = che; orange = par; purple = pil; yellow = SAM; red = Tn5042 mercury resistance transposon; pink = uvr; blue = tra. Loci are numbered according to the locus_tag feature.

**Supplementary Figure S5.**
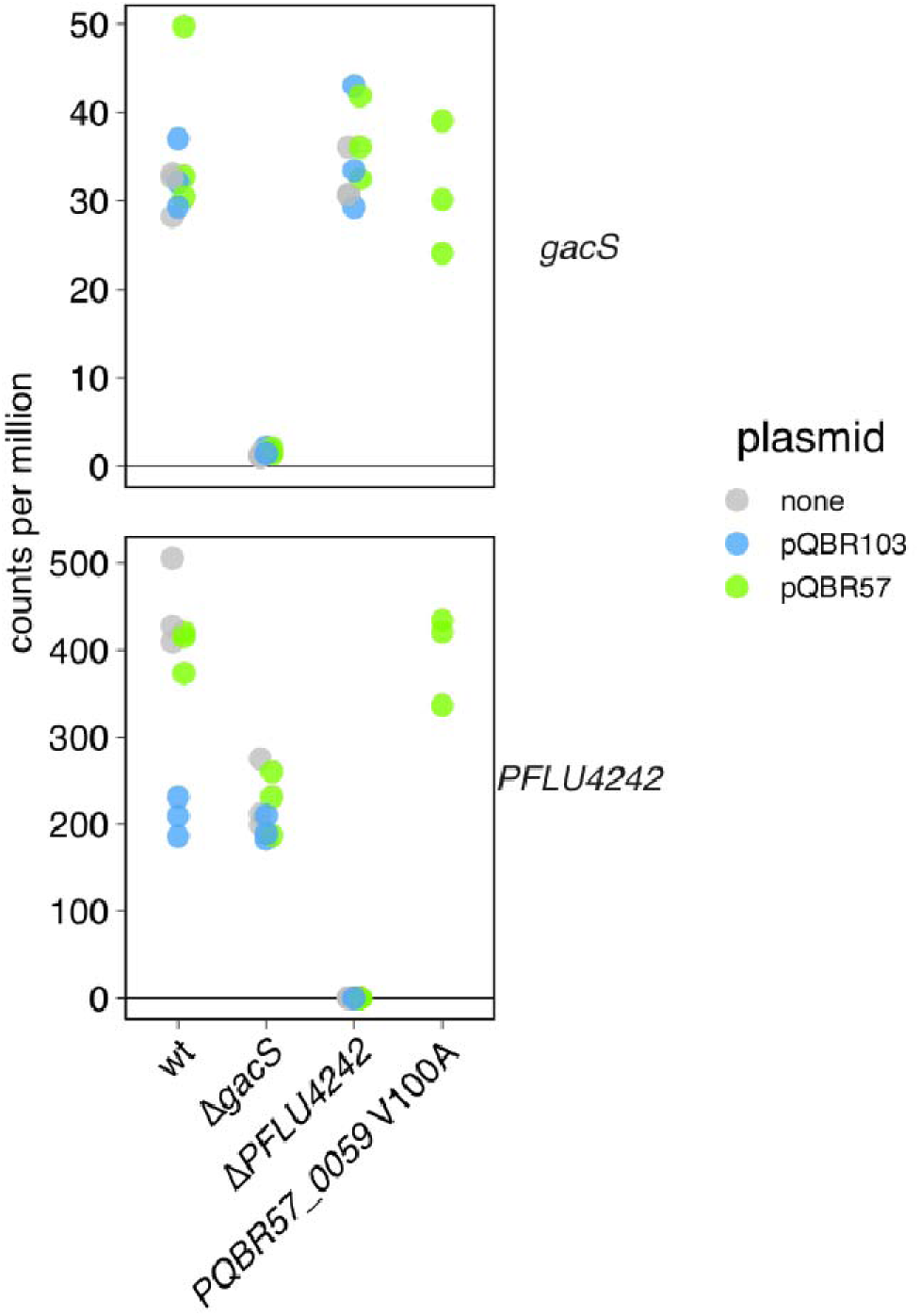
Expression of PFLU3777 (gacS) and PFLU4242 in the different treatments. Note different y-axis scaling. Points indicate different replicates, with values for each replicate scaled by effective library size.

**Supplementary Figure S6.**
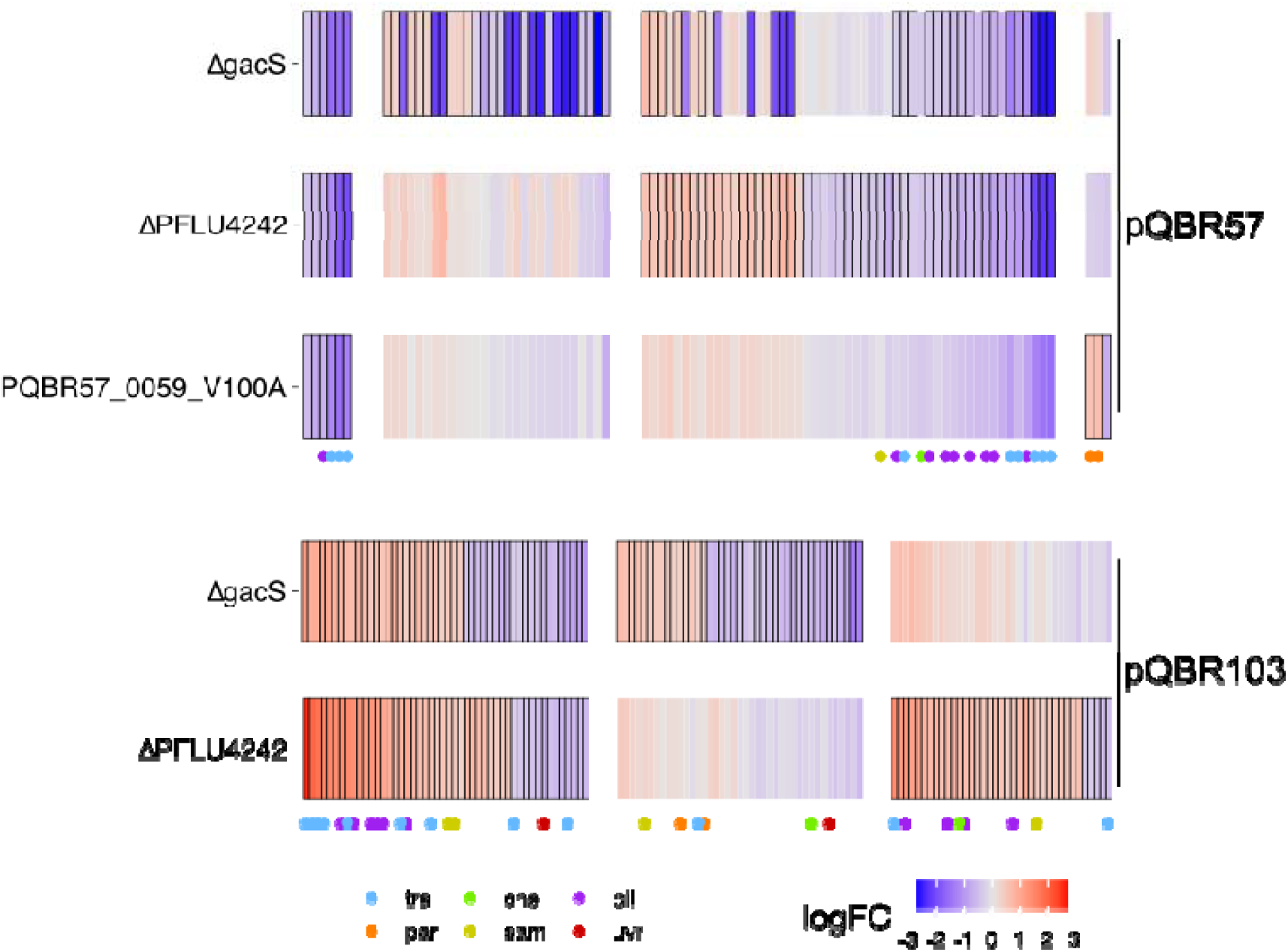
Differentially-expressed plasmid genes across all amelioration conditions. Image as **Fig. 4A**, but showing also the effect of ΔgacS and differentially-expressed genes on pQBR103.

